# *Pseudomonas aeruginosa* lectins LecA and LecB differentially engage and activate the EphA2 receptor

**DOI:** 10.64898/2026.11.24.768789

**Authors:** Carmen Senin, Olga Makshakova, Francois Naegele, Anselm Storbeck, Francesca Rosato, Maokai Xu, Pablo Rios-Munoz, Winfried Römer

## Abstract

*Pseudomonas aeruginosa* is a critical nosocomial pathogen responsible for severe and often difficult-to-treat infections in humans. Despite their central role in early host–pathogen interactions, the carbohydrate-binding lectins LecA and LecB remain largely underexplored compared to other *P. aeruginosa* virulence factors. Here, we show that LecA and LecB activate the human receptor tyrosine kinase EphA2 in a time-and carbohydrate-dependent manner in lung epithelial cells. Whereas LecB directly engages EphA2 primarily through protein–glycan binding, LecA activates EphA2 indirectly via interactions with the glycosphingolipid Gb3. Both lectins strongly colocalize with EphA2 at the host plasma membrane and trigger downstream CrkII phosphorylation via canonical EphA2 signaling. Owing to their tetravalent nature, LecA and LecB can cross-link membrane receptors, driving lateral membrane reorganization that initiates downstream signaling. Moreover, simultaneous engagement of both lectins at the plasma membrane further potentiates EphA2 activation. Finally, we demonstrate that live *P. aeruginosa* preferentially adheres to membrane domains where EphA2 concomitantly accumulates, inducing receptor activation in host cells. Collectively, these findings identify EphA2 as a pivotal host receptor mediating LecA-and LecB-driven signal transduction and provide new mechanistic insights into how *P. aeruginosa* exploits human lung epithelial cells during infection.

## Background

The Gram-negative, opportunistic bacterial pathogen *P. aeruginosa* poses a substantial global health challenge. Owing to its high metabolic versatility and adaptability to changing environments, it exhibits resistance to almost all clinically relevant antibiotics, leading to high morbidity and mortality in humans (1–4). Its diverse repertoire of virulence factors enables *P. aeruginosa* to invade host cells and subvert host physiology, often causing tissue damage and chronic infections, particularly in cystic fibrosis patients (5–8). Among these virulence factors are the homotetrameric lectins LecA (PAIL) and LecB (PAIIL), which localize to the bacterial cytoplasm, as well as the inner and outer membranes (9,10).

Both lectins possess high binding affinity/avidity for specific glycan structures. LecA preferentially binds α-D-galactose, a terminal carbohydrate motif present on its primary host receptor, the glycosphingolipid (GSL) globotriaosylceramide, also known as Gb3, CD77 or P^k^ blood group antigen. In contrast, LecB predominantly recognizes L-fucose and, with lesser affinity, D-mannose (11–14). Because of their distinct carbohydrate specificities, LecA and LecB modulate host cellular processes by engaging different surface receptors. For example, in human non-small cell lung cancer cells (H1299), LecA binding to Gb3 induces negative membrane curvature, promoting bacterial engulfment via the “lipid zipper” mechanism (15). Consistent with this role, a *P. aeruginosa* mutant lacking LecA (Δ*lecA*) shows significantly reduced invasiveness, establishing LecA as a primary driver of bacterial internalization. Downstream of receptor engagement, stimulation of H1299 cells with LecA triggers the Gb3-dependent phosphorylation of the adapter protein CT10 regulator of kinase II (CrkII) and Src family kinases (16). Furthermore, LecA–Gb3 interactions promote the recruitment of the glycosylphosphatidylinositol (GPI)-anchored protein CD59 to the outer leaflet and phosphatidylinositol (3,4,5)-trisphosphate (PIP₃) to the inner leaflet of newly formed membrane nanodomains (17), underscoring the critical role of these lipid-protein assemblies in *P. aeruginosa* host invasion.

In contrast, LecB binds to β1-integrins on the basolateral surface of polarized Madin-Darby canine kidney (MDCK) II cells, inducing integrin internalization and impairing cell migration and wound closure (18), which may facilitate *P. aeruginosa* colonization of injured epithelia (19,20). Moreover, LecB binds and cross-links fucosylated receptors on the apical membrane of polarized MDCK II cells, activating the Src-PI3K/AKT signaling pathway and driving the accumulation of PIP₃ clusters at the apical side (21). Despite these insights, the precise mechanism by which LecA and LecB engagement at the host plasma membrane is transduced into intracellular signaling cascades remains poorly understood. To address this question, we previously screened for potential host interaction partners, including membrane receptors, using a kinase assay and affinity pull-down coupled to mass spectrometry (22). Notably, members of the erythropoietin-producing hepatocellular (Eph) receptor family emerged among the most prominent candidates identified.

Eph receptors constitute the largest family of receptor tyrosine kinases (RTKs) in humans and are divided into EphA and EphB subfamilies. The EphA subfamily comprises nine receptors, which primarily interact with five GPI-anchored ephrin-A ligands, whereas the EphB subfamily includes five receptors and three transmembrane ephrin-B ligands. Cross-class interactions may occur. Members of both subfamilies share a conserved modular domain architecture consisting of an extracellular region, which contains a ligand-binding domain (LBD), a Sushi domain, a cysteine-rich EGF-like domain, and two fibronectin type III domains (FN1 and FN2), followed by a single transmembrane domain (TMD), an intracellular juxtamembrane (JM) region, a kinase domain, a C-terminal sterile alpha motif (SAM) and a PSD-95/Dlg1/ZO-1 (PDZ)-binding motif (23). Importantly, the ectodomain of EphA2 carries N-linked glycans, within its FN1 and FN2 domains (24,25).

Canonical Eph-ephrin signaling typically requires direct cell-cell contact, although it can also occur between non-contacting cells via extracellular vesicles such as exosomes that carry ligands or receptors (26). This canonical pathway involves receptor oligomerization and autophosphorylation of conserved tyrosine residues, including the two juxtamembrane residues Tyr588 and Tyr594, as well as tyrosine residues within the kinase domain (27–29). In EphA2, phosphorylation of Tyr588 relieves autoinhibition and generates docking sites for intracellular SH2 domain-containing signaling proteins, such as GEFs (28,30). Conversely, non-canonical signaling is ligand-independent and relies on serine/threonine phosphorylation (e.g., at Ser897) within the kinase-SAM linker region, mediated by kinases such as PKA or PKC, RSK, AKT, CK1, or Src (23,31).

Eph-ephrin signaling can proceed via forward (receptor-dependent), reverse (ligand-dependent), or bidirectional pathways. These cascades regulate diverse physiological and pathological processes across multiple cell types (including neurons, immune cells, stem cells, endothelial cells, and epithelial cells), ranging from axon guidance and synaptic plasticity to angiogenesis and cancer progression (31,32). Notably, Eph signaling, and EphA2 in particular, is also frequently co-opted during host–pathogen interactions.

Numerous studies have demonstrated that both viruses and bacteria exploit the EphA2 receptor to facilitate host cell infection (33–39). For instance, *Staphylococcus aureus* infections elevate EphA2 protein abundance and phosphorylation levels, whereas *EPHA2* knockout in HEK293T cells markedly reduces intracellular bacterial burden (40), establishing EphA2 as a key invasion factor. However, its role during *P. aeruginosa* infection remains unexplored. Here, we investigated whether *P. aeruginosa* lectins LecA and LecB, as well as live bacteria, interact with EphA2. Specifically, we examined whether these interactions depend on protein–carbohydrate recognition and trigger downstream CrkII signaling potentially to promote cytoskeletal remodeling and bacterial invasion.

Using complementary biochemical, microbiological and imaging approaches, this study demonstrates that LecA, LecB, and live *P. aeruginosa* induce time-dependent EphA2 phosphorylation, with LecA eliciting earlier and stronger receptor activation. Furthermore, we show that EphA2 activation depends on lectin–carbohydrate recognition, mediated primarily through the direct binding of LecB to N-linked glycans on EphA2 and indirect engagement of EphA2 by LecA via Gb3 binding. In addition, both lectins trigger EphA2 kinase-dependent phosphorylation of CrkII. Importantly, *P. aeruginosa* adhesion to host cells strongly correlates with EphA2 enrichment at the cell plasma membrane, and pharmacological inhibition of lectin– carbohydrate interactions substantially attenuates bacteria-induced EphA2 activation. Collectively, these findings establish EphA2 as a central host target linking lectin–receptor engagement to downstream intracellular signaling in lung epithelial cells.

## Materials and Methods

### Cell line, bacterial strains and cultivation conditions

The human non-small cell lung cancer epithelial cell line NCI-H1299 (American Type Culture Collection, CRL-5803, Manassas, VA, USA) was used for all experiments. This cell line was selected to investigate canonical EphA2 signaling in a physiologically relevant pulmonary context, as *P. aeruginosa* is a clinically important respiratory pathogen (41) and H1299 cells exhibit high endogenous EphA2 expression levels among non-small cell lung cancer cell lines (42–44). Cells were maintained in Roswell Park Memorial Institute (RPMI) 1640 medium, supplemented with 10% (v/v) fetal calf serum (FCS; Gibco, Thermo Fisher Scientific, Waltham, MA, USA) and 2 mM L-glutamine (Gibco, Thermo Fisher Scientific) at 37 °C in a humidified atmosphere containing 5% CO_2_.

The *P. aeruginosa* wild-type strain PAO1 (PAO1-WT; kindly provided by Prof. Dr. K.-E. Jäger, HHU Düsseldorf, Germany) was used for all infection assays and subsequent Western blot analyses. A PAO1 strain constitutively expressing mCherry (PAO1-mCherry; kindly provided by Prof. Dr. R. Kümmerli, University of Zurich, Switzerland) was used for immunofluorescence microscopy. All PAO1 strains were grown in Luria-Bertani (LB) broth (Carl Roth, Karlsruhe, Germany) for 16-18 h at 37 °C with continuous shaking at 180 rpm.

### Lectin production and stimulation with lectins and PAO1

Recombinant lectins LecA and LecB were expressed in the *Escherichia coli* strain BL21 (DE3), transformed with the expression vectors pET25-*pa1l* for LecA and pET25-*pa2l* for LecB (both kindly provided by Dr. A. Imberty, CNRS, Grenoble, France). The lectins were purified by affinity chromatography using D-galactose-agarose (Thermo Fisher Scientific, #20372, Schwerte, Germany) for LecA and D-mannose-agarose (Sigma Aldrich, #M6400, Taufkirchen, Germany) for LecB, as previously described by Chemani et al. (45). Following purification, LecA and LecB were diluted in 1× calcium-and magnesium-free phosphate-buffered saline (PBS⁻/⁻), sterile-filtered, and stored either at −80 °C for long-term storage or at 4 °C for up to 2 months.

For lectin stimulation assays, H1299 cells were seeded at 4.5 × 10^5^ cells/mL in 6-well culture plates (Merck, Taufkirchen, Germany). Prior to stimulation, cells were washed once with pre-warmed 1×

PBS⁻/⁻ to remove dead and non-adherent cells and serum-starved in FCS-free RPMI 1640 medium (starvation medium) for 2 h at 37 °C and 5% CO₂ to reduce baseline signaling. The medium was then replaced with starvation medium containing 300 nM LecA and/or LecB (calculated based on their homotetrameric molecular weight (MW)). Cells were pre-incubated on ice for 30 min to allow synchronized lectin binding to the plasma membrane without internalization, followed by a temperature shift to 37 °C and 5% CO_2_ for 15 min to 4 h without medium exchange.

For bacterial infection assays, H1299 cells (seeded at 4.5 × 10^5^ cells/mL) were cultured and serum-starved for 2 h at 37 °C and 5% CO_2_, and all subsequent steps were performed in starvation medium. Suspensions of PAO1-WT were adjusted to a multiplicity of infection (MOI) of 15, based on optical density measurements at 600 nm (OD₆₀₀). To inhibit host EphA2 kinase activity, H1299 cells were pre-treated with the type II small-molecule inhibitor ALW-II-41-27 (500 nM, MedChemExpress, #HY-18007, Hamburg, Germany) for 45 min prior to infection. Cells were then exposed to four bacterial treatment conditions: (1) untreated PAO1-WT; (2) PAO1-WT pre-incubated with ALW-II-41-27 (500 nM) for 45 min at room temperature (RT); (3) PAO1-WT pre-incubated with 10 mM *p*-nitrophenyl α-D-galactopyranoside (pNPG; Thermo Fisher Scientific, #L13376.ME, Germany) and 45 mM L-fucose (Sigma-Aldrich, #F2252, Germany) for 30 min at RT to saturate the lectin carbohydrate-binding pockets; and (4) PAO1-WT pre-incubated with all three compounds for 30 min at RT (30 min with pNPG and L-fucose, and 45 min with ALW-II-41-27). PBS⁻/⁻ served as the vehicle control throughout all experiments, as both pNPG and L-fucose were dissolved in this buffer. Following pre-incubation at RT, the bacterial suspensions were added to the cells and incubated for 2 h at 37 °C and 5% CO₂.

Cells and bacteria were subsequently lysed in ice-cold radio-immunoprecipitation assay (RIPA) buffer (20 mM Tris HCl [pH 8.0], 0.1% [w/v] SDS, 10% [v/v] glycerol, 13.7 mM NaCl, 2 mM EDTA, 0.5% [w/v] sodium deoxycholate), supplemented with protease and phosphatase inhibitors: 1 µM sodium orthovanadate, 0.2 µM pefablock and 1% (v/v) phosphatase inhibitor cocktail 3 (all from Merck, #567540-5GM, #76307, #P0044, Germany). Lysates were boiled in 5× Laemmli SDS-PAGE sample buffer for 15 min to inactivate bacteria and denature proteins prior Western blot analysis.

### Lectin fluorescence labelling and biotinylation

LecA and LecB were fluorescently labelled with Alexa Fluor (AF) 647, AF555, or Cyanine (Cy) 3 dyes (Thermo Fisher Scientific, Germany) via a *N*-hydroxysuccinimide (NHS) ester conjugation. Briefly, lectins dissolved in 1× PBS⁻/⁻ were supplemented with 1 M NaHCO_3_ (pH 8.5) at a 1:10 (v/v) ratio. The fluorophores were dissolved in DMSO (Thermo Fisher Scientific, Germany) at a concentration of 50 µg/mL and added to the protein solutions at a 6-fold molar excess. The reaction mixtures were incubated for 1 h at RT under continuous rotation, protected from light. Unreacted fluorophore was subsequently removed using Zeba Spin desalting columns (7K MWCO), Thermo Fisher Scientific, #89878, Germany). Labelled protein concentrations were determined using a NanoDrop 2000c photometer (Thermo Fisher Scientific, Germany).

For biotinylation, LecA and LecB were modified using the EZ-Link NHS-PEG4 Biotinylation Kit (Thermo Fisher Scientific, #21455, Germany) according to the manufacturer’s instructions. Briefly, EZ-Link NHS-PEG4-Biotin was dissolved in ultrapure water and added to each lectin solution at a 20-fold molar excess. Following incubation for 1 h at RT under continuous rotation, unbound biotin was removed using Zeba Spin desalting columns (7K MWCO). Biotinylation efficiency was determined using a 4′-hydroxyazobenzene-2-carboxylic acid (HABA) assay included in the kit.

### SDS-PAGE, Western blot and lectin blot

Lectin-stimulated H1299 cells were lysed in ice-cold RIPA buffer supplemented with sodium orthovanadate (1 µM), Pefablock (0.2 µM) and 1% (v/v) Phosphatase Inhibitor Cocktail 3. Cells were incubated for 20 min at 4 °C under agitation (150 rpm), scraped from the culture plates, and incubated for an additional 15 min under the same conditions. Lysates were subsequently collected, sonicated, and clarified by centrifugation at 21,000 × g for 15 min at 4 °C. Protein concentrations in the supernatants were determined using the Pierce bicinchoninic acid (BCA) protein assay (Thermo Fisher Scientific, #23227, Germany). Normalized protein samples were mixed with 5× Laemmli SDS-PAGE sample buffer and denatured at 98 °C for 8 min. Equal amounts of protein were resolved on 10% or 12% acrylamide gels and transferred onto nitrocellulose membranes (0.22 µm or 0.45 µm pore size) via wet electroblotting in a Tris-glycine buffer containing 20% (v/v) methanol (60 min at 400 mA per two gels). Membranes were blocked for 1 h at RT in 1× TBS-T containing either 3% (w/v) bovine serum albumin (BSA, Serva, Heidelberg, Germany) for phospho-protein detection or 2-5% (w/v) skim milk powder (Carl Roth, Karlsruhe, Germany) for total protein detection.

Membranes were incubated overnight at 4 °C under gentle agitation (20 rpm) with the following primary antibodies diluted in the respective blocking buffers: monoclonal anti-phospho-EphA2 (Tyr588, 1:500, Cell Signaling Technology, #12677, Leiden, The Netherlands), monoclonal anti-EphA2 (1:500, Cell Signaling Technology, #6997, The Netherlands), polyclonal anti-phospho-CrkII (Tyr221, 1:500, Cell Signaling Technology, #3491, The Netherlands), monoclonal anti-CrkII (1:500, BD Transduction Laboratories, #610035, Laagstraat, Belgium), and polyclonal anti-GAPDH (1:1,000, Merck, #G9545, Germany). The following day, membranes were washed three times in 1× TBS-T and incubated for 1 h at RT with HRP-conjugated secondary antibodies (anti-rabbit, #7074 or anti-mouse, #7076, 1:2,000, Cell Signaling Technology, The Netherlands). Protein bands were visualized using the Novex ECL Chemiluminescent Substrate Reagent Kit (Thermo Fisher Scientific, #WP20005, Germany), and imaged on a ChemiDoc MP Imaging System (Bio-Rad). Phospho-protein levels were quantified using the GelAnalyzer software (v23.1.1) and normalized to the total protein load determined by Ponceau S staining.

Prior to SDS-PAGE and lectin blotting, cell lysates were treated with Peptide:N-glycosidase F (PNGase F, Merck, #P7367-50UN, Germany) to remove N-linked oligosaccharides. To this end, 90 µL of cell lysate was mixed with 10 µL of denaturing solution (0.2% SDS, 100 mM β-mercaptoethanol) and boiled at 100 °C for 10 min. After cooling briefly on ice, Triton X-100 (15% [v/v], added at a 1:10 [v/v] ratio) and 3 µL of PNGase F per 100 µL of denatured lysate were added, followed by incubation at 37 °C for 2.5 h. Subsequently, 5× Laemmli SDS-PAGE sample buffer was added, and the samples were boiled again for 5 min to inactivate the PNGase F. Proteins were resolved on 10% or 12% polyacrylamide gels and transferred onto a nitrocellulose membrane (0.22 µm pore size) via wet electroblotting. Non-specific binding sites were blocked by incubating the membrane in Intercept Blocking Buffer (LICORbio, #927-90001, Bad Homburg, Germany). Fluorescently labelled LecA-AF647 (300 nM) and LecB-AF555 (1 µM, concentrations calculated based on the tetrameric MW) were diluted in blocking buffer and incubated with the membrane overnight at 4 °C with gentle agitation (20 rpm). Lectin binding was visualized using a ChemiDoc MP Imaging System applying the AF647 or Cy3 channels, respectively. Following lectin detection, the membrane was incubated overnight at 4 °C while gentle agitation (20 rpm) with a monoclonal anti-EphA2 primary antibody (Cell Signaling Technology, #6997, The Netherlands) diluted 1:500 in blocking buffer. After washing, an anti-rabbit AF488-conjugated secondary antibody (Thermo Fisher Scientific, Germany) was applied for 1 h at RT. Finally, the membrane was re-imaged, and signal co-localization was assessed by overlaying the AF647/AF555 and AF488 channels.

### Inhibitor studies

To saturate the carbohydrate-binding pockets of LecA and LecB, pNPG and L-fucose were used, respectively. To deplete cells of glucosylceramide (GlcCer)-based glycosphingolipids (GSLs), the GlcCer synthase inhibitor D-*threo*-1-phenyl-2-palmitoylamino-3-morpholino-1-propanol (PPMP, Santa Cruz Biotechnology, #sc-205655, Dallas, TX, USA) was applied. For pNPG and L-fucose inhibition assays, and subsequent lectin stimulations, cells were seeded at a density of 4.5 × 10^5^ cells/mL, and incubated in serum-free (starvation) medium for 2 h at 37 °C and 5% CO_2_. All subsequent experimental steps were performed in starvation medium. Prior to stimulation, LecA or LecB was pre-incubated with pNPG (10 mM) or L-fucose (45 mM), respectively, for 30 min at RT. Next, the lectins (300 nM final concentration) were added to the cells in presence or absence of their respective inhibitory carbohydrates. Cells were initially incubated on ice for 30 min to synchronize lectin binding and subsequently transferred to a humidified incubator (37 °C, 5% CO₂) for 1 h or 2 h without medium exchange.

For glycosphingolipid depletion, cells were treated with 2.5 µM PPMP for 4 d until reaching a density of 4.5 × 10⁵ cells/mL. Prior to lectin stimulation, cells were incubated in starvation medium for 2 h at 37 °C and 5% CO_2_. Subsequently, cells were stimulated with 300 nM LecA or LecB in the presence of PPMP or left unstimulated with PPMP alone. DMSO (0.013% [v/v]) served as the vehicle control. Cells were initially incubated on ice for 30 min to allow lectin binding, and then transferred to a humidified incubator (37 °C, 5% CO₂) for 1 h or 2 h without medium exchange.

To inhibit EphA2 receptor kinase activity, the ATP-competitive tyrosine kinase inhibitor ALW-II-41-27 was used. Cells (4.5 × 10⁵ cells/mL) were serum-starved for 2 h at 37 °C and 5% CO₂, followed by pre-incubation with 500 nM ALW-II-41-27 for 45 min at 37 °C. Immediately following pre-incubation, fresh inhibitor (500 nM) was added in the presence or absence of 300 nM LecA or LecB, with DMSO (0.5% [v/v]) serving as the vehicle control. Cells were incubated on ice for 30 min and subsequently shifted to 37 °C (5% CO₂) for 15 min to 2 h without medium exchange.

To assess LecA binding to PPMP-treated cells via immunofluorescence, cells were seeded at a density of 1 × 10^4^ cells/mL onto 12 mm glass coverslips, allowed to adhere overnight, and treated with 2.5 µM PPMP for 4 d. Following PPMP pre-treatment, cells were incubated with either fluorescently labelled LecA-AF647 (300 nM) or the Gb3-specific and fluorescently labelled Shiga toxin B-subunit (StxB-AF647, 260 nM based on the pentameric MW). Cells were initially incubated on ice for 30 min to allow lectin binding and subsequently transferred to a humidified incubator (37 °C, 5% CO₂) for 1 h without medium exchange. Next, cells were fixed with ice-cold 4% PFA for 15 min, and excess aldehyde groups were quenched with 50 mM NH_4_Cl for 10 min at RT. To block non-specific binding and permeabilize the cell membrane, cells were incubated in blocking/permeabilization buffer consisting of 3% BSA and 0.2% saponin in 1× PBS⁻/⁻ for 35 min at RT. Filamentous actin (F-actin) and nuclei were counterstained using Atto 565-conjugated Phalloidin (1:1,000, ATTO-TEC, #AD565-81, Siegen, Germany) and DAPI (1:500, Carl Roth, #6335.1, Germany), respectively. Finally, stained coverslips were mounted onto glass microscope slides using 8 µL of Mowiol supplemented with the antifade reagent 1,4-diazabicyclo [2.2.2] octane (DABCO, Sigma-Aldrich, #D27082, Germany).

### Immunofluorescence experiments and bacterial imaging

H1299 cells were seeded at a density of 1.5 × 10^5^ cells/mL onto 12 mm glass coverslips and allowed to adhere overnight. Fluorescently labelled LecA-AF647 and LecB-Cy3 were added to the cells either simultaneously at equimolar concentrations (300 nM each) or individually at equimolar concentrations (300 nM) or at distinct concentrations (300 nM LecA, 1 µM LecB). Cells were initially incubated on ice for 30 min to allow lectin binding and subsequently transferred to a humidified incubator (37 °C, 5% CO₂) for 15 min to 2 h without medium exchange. Next, cells were fixed with ice-cold 4% PFA for 15 min, and excess aldehyde groups were quenched with 50 mM NH₄Cl for 10 min at RT. Cells were then blocked and permeabilized in 3% BSA and 0.2% saponin in 1× PBS⁻/⁻ for 35 min at RT. Subsequently, samples were incubated for 1 h at RT with the following primary antibodies: monoclonal anti-EphA2 (1:100, Cell Signaling Technology, #6997, The Netherlands) and monoclonal anti-Giantin (cis/medial Golgi marker, 1:100, Abcam, #ab37266, Amsterdam, The Netherlands). Following washing, cells were incubated for 30 min at RT with DyLight 488-conjugated anti-rabbit (1:200, Thermo Fisher Scientific, #35552, Germany) or Cy3-conjugated anti-rabbit (1:200; Merck, #AP182C, Germany) secondary antibodies, supplemented with DAPI (1:500) for nuclear counterstaining. Finally, coverslips were mounted onto glass microscope slides.

For *P. aeruginosa* and EphA2 localization analysis, a *P. aeruginosa* PAO1 strain constitutively expressing mCherry was used. Based on OD₆₀₀ measurements, bacteria were added at a MOI of 15 to the cells (seeded at 1 × 10⁵ cells/mL) grown onto 12 mm glass coverslips at 37 °C and 5% CO₂. Following co-incubation for 2 h at 37 °C and 5% CO₂, infected cells were fixed with ice-cold 16% PFA for 20 min at RT, and excess aldehyde groups were quenched with 50 mM NH₄Cl for 10 min at RT. Cells were then blocked and permeabilized in 3% BSA and 0.2% saponin in 1× PBS⁻/⁻ for 35 min at RT. For EphA2 immunostaining, samples were incubated with a monoclonal anti-EphA2 primary antibody (1:100; Cell Signaling Technology, #6997, The Netherlands) followed by a DyLight 488-conjugated anti-rabbit secondary antibody (1:200; Thermo Fisher Scientific, #35552, Germany). Nuclei and bacterial DNA were counterstained with DAPI (1:500).

All immunofluorescence samples were imaged using a confocal laser scanning microscope (Nikon A1R) equipped with a 60× oil-immersion objective (numerical aperture [NA] = 1.49). Colocalization analysis of LecA-AF647, LecB-Cy3 (or LecB-AF647), and EphA2-AF488 at the plasma membrane was performed in Fiji (ImageJ v2.16.0/1.54p, Java 1.8.0_172 [64-bit], National Institutes of Health, USA) using the BIOP-JACoP plugin with Costes’ automatic thresholding. To isolate plasma membrane signals, regions of interest (ROIs) were manually delineated to exclude intracellular fluorescence across individual focal planes of each z-stack. Only ROIs yielding thresholded Manders’ colocalization coefficients (M1 and M2) indicative of low background noise were included in the final analysis. To visualize bacterium–host cell membrane interactions, five sequential z-planes from both infected or control samples were combined as maximum-intensity projections (or z-projections) to generate pseudo-3D composite images.

### Pull-down assay and immunoprecipitation

For lectin pull-down assays, biotinylated LecA or LecB was added to the cells (4.5 × 10⁵ cells/mL) at equimolar concentrations (300 nM). Cells were initially incubated on ice for 30 min to allow lectin binding and subsequently transferred to a humidified incubator (37 °C, 5% CO₂) for 1 h or 3 h without medium exchange. Cells were then lysed in ice-cold pull-down lysis buffer (25 mM Tris-HCl [pH 7.4], 150 mM NaCl, 1 mM EDTA, 1% [v/v] IGEPAL CA-630 [NP-40], and 5% [v/v] glycerol) supplemented with 11 µM leupeptin (Merck, #L9783, Germany), 0.2 µM Pefabloc, and 1% (v/v) Phosphatase Inhibitor Cocktail 3. Lysates were cleared by centrifugation at 21,000 × g for 15 min at 4 °C, and a 100 µL aliquot of the cleared supernatant was retained as the input control. The remaining supernatant was adjusted to a final volume of 300 µL with binding/wash buffer (0.1% [v/v] Tween-20 in TBS) and incubated with 75 µL of pre-washed Pierce Streptavidin Magnetic Beads (Thermo Fisher Scientific, Germany) for 1 h at RT with end-over-end rotation. Following magnetic separation, the supernatant was collected as the flow-through fraction. Bound proteins were eluted from the beads in 1× Laemmli SDS-PAGE sample buffer, magnetically separated, and analyzed by Western blotting.

For co-immunoprecipitation of EphA2 or β1-integrin from unstimulated H1299 cells, whole-cell lysates were prepared using ice-cold IGEPAL lysis buffer (50 mM Tris-HCl [pH 7.4], 150 mM NaCl, 1 mM EDTA, and 1% [v/v] IGEPAL CA-630) supplemented with 11 µM leupeptin, 0.2 µM Pefabloc, and 1% (v/v) Phosphatase Inhibitor Cocktail 3. After retaining an unprocessed aliquot as the input control, the remaining lysate was pre-cleared with 20–50 µL of superparamagnetic Protein G Dynabeads resuspended in 1× PBS⁻/⁻ for 20 min at RT with gentle rotation. The pre-cleared lysate was then incubated overnight at 4 °C with rotation in the presence of monoclonal anti-EphA2 (1:100, Cell Signaling Technology, #6997, The Netherlands) or monoclonal anti-β1-integrin (1:100; Thermo Fisher Scientific, #MA1-19105, Germany) primary antibodies. The following day, 200–500 µL of the antibody-lysate mixture was incubated with 20–50 µL of pre-washed Protein G Dynabeads for 20 min at RT with rotation. After magnetic separation, the supernatant was collected as the flow-through fraction. Immunoprecipitated complexes were eluted by boiling the beads in 1× Laemmli SDS-PAGE sample buffer at 95 °C for 7 min. All fractions (input, flow-through, and eluate) were analyzed by Western blotting using anti-EphA2 (1:500) or anti-β1-integrin (1:1,000) primary antibodies.

### Protein docking and molecular dynamics simulations

The intermolecular interaction modes between the ectodomain of EphA2 and the *P. aeruginosa* lectins LecA and LecB were investigated using *in silico* approaches. Crystal structures of the EphA2 ectodomain, LecA, and LecB were retrieved from the Protein Data Bank (PDB codes: 3FL7, 5D21 and 5A3O, respectively) and refined by adding missing loops with Modeller v9.15 (46). Protein–protein docking of the EphA2 ectodomain (residues 1-507) against each tetrameric lectin was performed using the ClusPro web server (47). The top ten docking poses were selected for further analysis, excluding redundant poses generated by rotational symmetry of the lectin homotetramers. The most plausible intermolecular complexes predicted by docking (four for LecB and two for LecA) were subjected to molecular dynamics (MD) simulations using Gromacs2025.0 (48,49) with the Charmm36m force-field (50). Each protein complex was solvated in a TIP3P water box under periodic boundary conditions, and Na^+^ ions were added to neutralize the net system charge. Equations of motion were integrated using a 2 fs time step, with all covalent bonds involving hydrogen atoms constrained via the LINear Constraint Solver (LINCS) algorithm. Long-rang electrostatic interactions were calculated using the Particle Mesh Ewald (PME) method. Simulations were conducted in the isothermal-isobaric (*NPT*) ensemble at 300 K using a velocity-rescaling (V-rescale) thermostat (coupling time constant = 0.1 ps) and a Parrinello-Rahman barostat (relaxation time = 2 ps; isothermal compressibility = 4.5×10^-5^ bar^-1^). Following energy minimization and equilibration, a 500 ns production run was performed for each complex, with the final 200 ns used to calculate binding free energies via the molecular mechanics/generalized Born surface area (MM/GBSA) method. Complexes that maintained stable protein–protein contacts and reached a root-mean-square deviation (RMSD) plateau over the 500 ns trajectory were selected as plausible interaction models for EphA2 and each lectin.

### Statistics

All data were analyzed and visualized using GraphPad Prism v10.8.2 (GraphPad Software, San Diego, CA, USA). Data are presented as the mean ± standard deviation (SD) from at least *n* = 3 independent biological replicates per condition. Multi-group comparisons across time points were performed using two-way analysis of variance (ANOVA) with treatment and time as independent factors. When significant main effects or interactions were detected, Šidák’s multiple-comparison *post hoc* test was applied to compare treatment groups against the corresponding control group at each time point, as well as to perform pairwise comparisons among treatment groups while controlling the family-wise error rate. Differences between two groups at a single time point were assessed using unpaired two-tailed Student’s *t*-tests. In all analyses, a *p*-value < 0.05 was considered statistically significant, with significance levels denoted by asterisks in the figures.

## Results

### LecA and LecB induce sustained EphA2 activation through carbohydrate binding

To determine how *P. aeruginosa* lectin binding to the host plasma membrane triggers intracellular signaling, we analyzed EphA2 activation in H1299 cells following LecA and LecB stimulation. To this end, we quantified Tyr588 phosphorylation using Western blotting. Compared to the PBS control, both LecA stimulation at 30 min (p = 0.0324) and LecB stimulation at 1 h (p = 0.0229) significantly increased EphA2 activation (p-EphA2). Phosphorylation levels peaked at 2 h of stimulation with each lectin, with a 3-fold (LecA) and 2.5-fold (LecB) increase relative to the PBS control, followed by a decline at 4 h (Fig. 1A, B). While LecA triggered faster and stronger EphA2 activation than LecB alone, co-stimulation with both lectins amplified this response, driving a nearly 4-fold increase in p-EphA2 levels at 2 h. No further increase in activation occurred after 4 h of co-stimulation, which may reflect saturation of lectin binding at the plasma membrane. Collectively, these data indicate an additive effect of LecA and LecB on EphA2, resulting in stronger and more sustained activation than stimulation with either lectin alone. Strikingly, these slow-onset kinetics contrast with the rapid activation (30 s to 10 min) typically induced by soluble ephrin-A1-Fc or endogenous ephrin-A1, which rapidly recruits downstream effectors such as SHP2, FAK, and ERK (51–54). Based on these findings, we propose that lectin-induced EphA2 activation, while distinct from classical ligand-dependent signaling, occurs through a mechanism that relies on receptor phosphorylation at Tyr588.

**Fig. 1.**
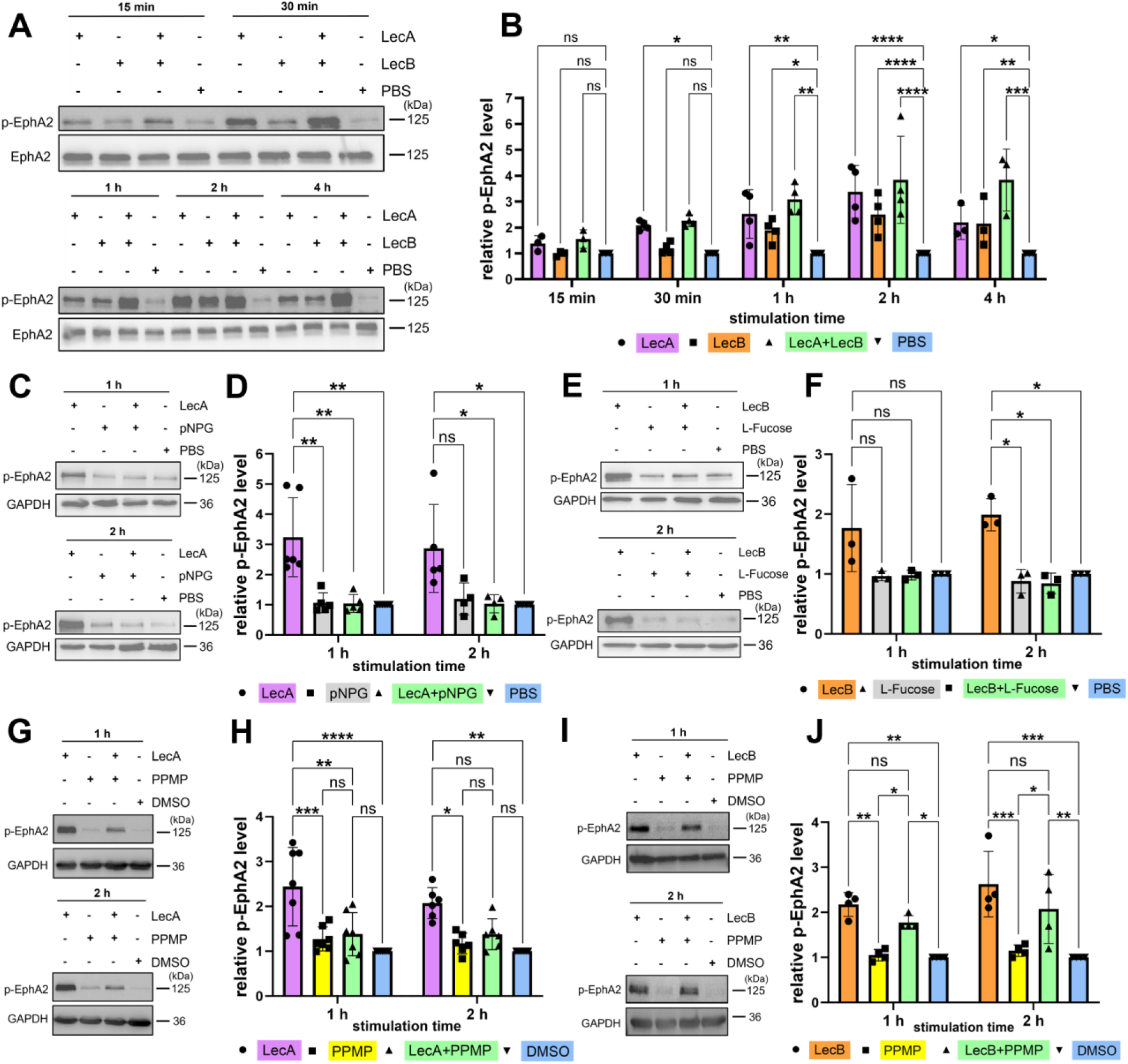
LecA and LecB differentially trigger EphA2 activation in a carbohydrate-dependent manner. **(A)** Representative Western blot images showing p-EphA2 levels following stimulation with LecA and LecB, which were applied to H1299 cells at equimolar concentrations (300 nM, based on the tetrameric MW) for 15 min to 4 h at 37 ℃. Total (pan-) EphA2 levels remained unchanged across treatments. PBS served as a control. **(B)** Quantification and statistical analysis of p-EphA2 levels from independent Western blot experiments, including those shown in (A). **(C) - (J)** H1299 cells were incubated for 1 h or 2 h at 37 °C with pNPG (10 mM), L-fucose (45 mM), or PPMP (2.5 µM) in the presence or absence of LecA or LecB (300 nM), alongside controls treated with LecA or LecB alone. GAPDH served as loading control and PBS or DMSO served as a treatment control. **(C)** and **(E)** Representative Western blot images illustrate a strong reduction of p-EphA2 levels after incubation of LecA and LecB with pNPG or L-fucose, respectively. **(D)** and **(F)** Quantification and statistical analysis of p-EphA2 levels from independent Western blot experiments, including those shown in (C) and (E). **(G)** and **(I)** Representative Western blot images illustrate a stronger reduction of LecA-induced p-EphA2 levels after PPMP treatment in contrast to LecB-induced p-EphA2 levels. **(H)** and **(J)** Quantification and statistical analysis of p-EphA2 levels across independent Western blot replicates, including those shown in (G) and (I). Prior to statistical analysis, densitometric values were normalized to the total protein load via Ponceau S staining and expressed relative to the respective PBS or DMSO control. Data are presented as mean ± SD from biological replicates (*n* ≥ 3 per group for pNPG and L-fucose treatments; *n* ≥ 4 per group for lectin stimulations without inhibitor and with PPMP treatment). Statistical significance was determined by ordinary two-way ANOVA followed by Šídák’s multiple comparisons test (ns, not significant; *p < 0.05; **p < 0.01; ***p < 0.001; ****p < 0.0001).

Despite the inherently weaker affinity of monomeric carbohydrate–lectin interactions, the high multivalent avidity of LecA and LecB toward α-galactosylated and fucosylated targets, respectively, was shown to be crucial for host receptor binding and subsequent bacterial internalization. For instance, applying galactose-based LecA inhibitors significantly diminishes *P. aeruginosa* host cell invasion. While monovalent pNPG competitively inhibits LecA binding to host cell receptors and lowers bacterial internalization by up to 60% (55), a divalent galactose ligand (0.05–5 µM) reduces H1299 cell invasion by up to 90% (56). Furthermore, L-fucose treatment was shown to block LecB-mediated integrin endocytosis, thereby preventing the associated impairment of MDCK II cell migration and wound healing (18). Therefore, we investigated whether the simultaneous treatment of H1299 cells with lectins and competitive inhibitors for 1 h or 2 h, corresponding to the observed maximum of EphA2 phosphorylation (Fig. 1A, B), affects lectin-induced EphA2 activation. Stimulation with LecA or LecB in the presence of pNPG or L-fucose, respectively, drastically reduced p-EphA2 levels back to PBS control levels (Fig. 1C–F; almost 100% reduction), indicating that lectin–carbohydrate interactions are essential for EphA2 activation.

Next, to determine whether other plasma membrane components are required for lectin-induced EphA2 activation, we depleted GlcCer-based GSLs (including Gb3, the main LecA receptor in H1299 cells) by PPMP treatment and examined subsequent EphA2 phosphorylation. In parallel, we verified the depletion of Gb3 by plasma membrane binding of the fluorescently labelled B-subunit of Shiga toxin (StxB). Consistent with successful Gb3 depletion, StxB exhibited virtually no binding to PPMP-treated cells, whereas LecA retained minor residual binding (Supplementary Fig. 1). Analysis of p-EphA2 levels revealed that LecA stimulation of PPMP-treated cells led to a marked reduction in phosphorylation compared to unstimulated DMSO-treated control cells (Fig. 1G, H; ∼75% reduction). Remarkably, these p-EphA2 levels remained slightly elevated relative to untreated controls, suggesting weak LecA-mediated EphA2 activation through a GSL-independent mode of receptor engagement. In contrast, LecB stimulation of PPMP-treated cells resulted in a substantially smaller decrease in p-EphA2 levels (Fig. 1J; ∼30% reduction compared to DMSO-treated control cells). Collectively, these data demonstrate that while GlcCer-based GSLs are critical for LecA-induced EphA2 activation, they play a much smaller role in that of LecB.

### Lectins regulate CrkII activity in an EphA2 kinase-dependent manner

Previous studies have demonstrated that LecA induces phosphorylation of CrkII at Tyr221 in H1299 cells (16). Building on these findings, we examined Tyr221 phosphorylation in cells with intact or impaired EphA2 kinase activity to further assess canonical downstream signaling upon lectin stimulation.

Treatment of H1299 cells with the Eph kinase inhibitor ALW-II-41-27 markedly reduced basal EphA2 phosphorylation, even below control levels (Fig. 2A, B). Furthermore, ALW-II-41-27 potently blocked EphA2 signaling, as simultaneous stimulation with LecA or LecB elicited only a marginal increase in p-EphA2.

**Fig. 2.**
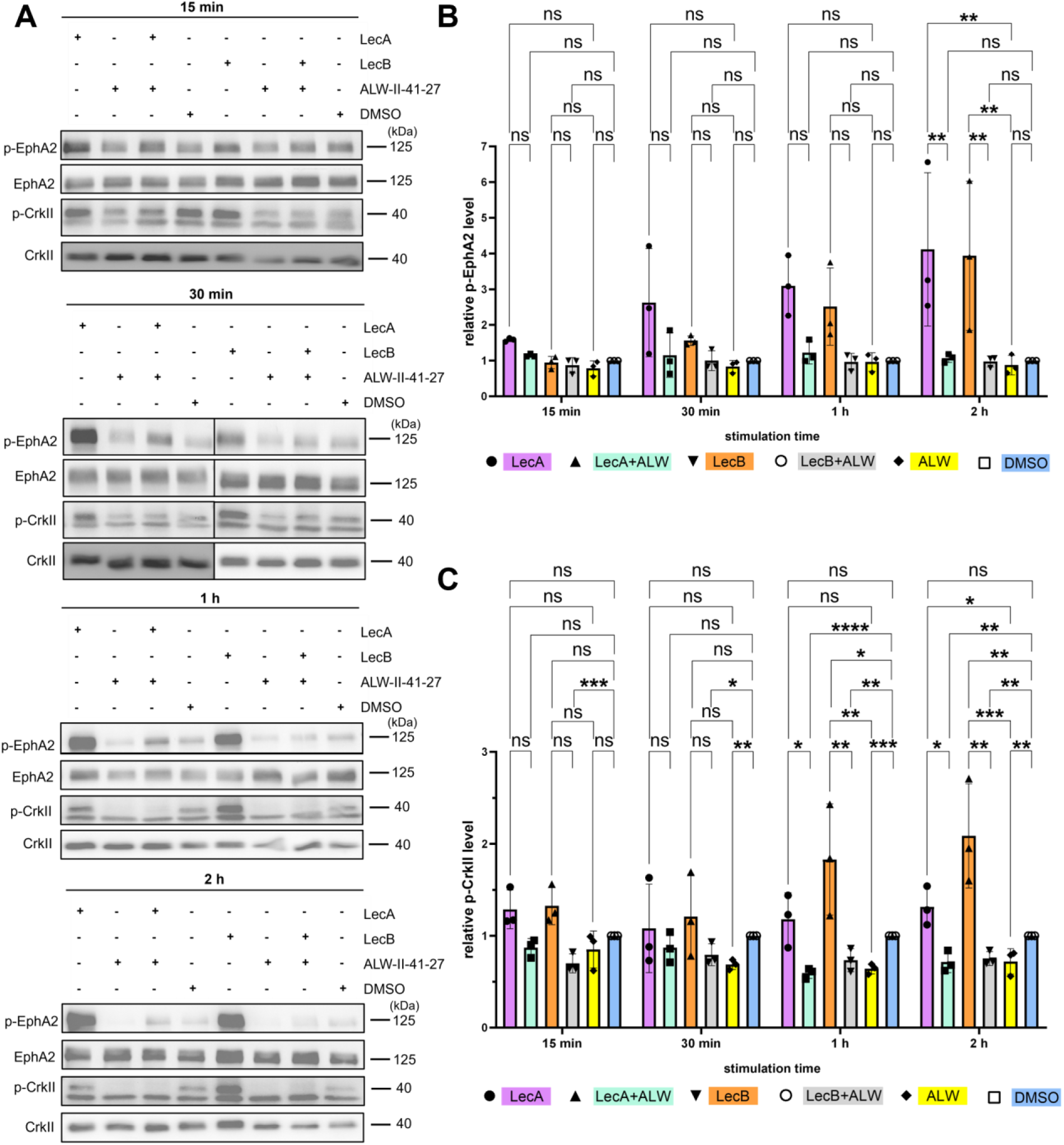
**LecA-and LecB-induced CrkII phosphorylation requires EphA2 kinase activity**. **(A)** Representative Western blot images showing p-EphA2 and p-CrkII levels in H1299 cells following treatment with ALW-II-41-27 and lectin stimulation. Cells were either left untreated or pre-treated with 500 nM ALW-II-41-27 for 45 min at 37 °C. Subsequently, cells were stimulated with equimolar concentrations (300 nM) of LecA or LecB for 15 min to 2 h at 37 °C, either alone or in the presence of freshly added ALW-II-41-27 (LecA + ALW, LecB + ALW). Cells treated with freshly added inhibitor alone (ALW) and DMSO-treated cells served as controls. Total (pan-) EphA2 levels remained unchanged across all treatment conditions. **(B, C)** Quantification and statistical analysis of p-EphA2 (B) and p-CrkII (C) levels from independent Western blot experiments, including the representative blots shown in (A). Densitometric values were normalized to total protein load via Ponceau S staining and expressed relative to the DMSO control. Data are presented as mean ± SD of *n* ≥ 3 biological replicates per group. Statistical analysis was performed using ordinary two-way ANOVA followed by Šídák’s multiple comparisons test (ns, not significant; *p < 0.05; **p < 0.01; ***p < 0.001; ****p < 0.0001).

Upon LecB stimulation, we observed a time-dependent increase in CrkII phosphorylation at Tyr221 (p-CrkII) that peaked at 2-fold relative to the DMSO control (Fig. 2A, C; 2 h). Conversely, LecA stimulation elicited a weaker p-CrkII response (1.3-fold relative to the DMSO control). These findings highlight a discrepancy between the activation of EphA2 and its downstream effector CrkII: while LecA induced earlier and stronger EphA2 phosphorylation, LecB triggered substantially higher p-CrkII levels. This suggests that the two lectins exploit distinct modes of EphA2 engagement and downstream signaling modulation. Crucially, ALW-II-41-27 treatment almost completely abolished CrkII phosphorylation (Fig. 2A, C), demonstrating that lectin-induced p-CrkII strictly depends on EphA2 kinase activity.

### LecA and LecB binding strongly correlates with plasma membrane-localized EphA2

Given that LecA-and LecB-induced EphA2 activation depends strongly on carbohydrate binding, with GSLs playing an essential role particularly for LecA, we next examined via immunofluorescence microscopy whether both lectins colocalize with EphA2 within the same plasma membrane domains upon host cell binding.

To this end, H1299 cells were treated either simultaneously (Fig. 3A) or individually (Supplementary Fig. 2) with fluorescently labeled LecA (AF647, magenta) and LecB (Cy3, yellow/magenta). Following immunofluorescence staining of EphA2 (AF488, cyan), colocalization was quantified using Manders’ colocalization coefficients (MCCs), where M1 and M2 denote the fractions of lectin overlapping with EphA2 and EphA2 overlapping with lectin, respectively.

**Fig. 3.**
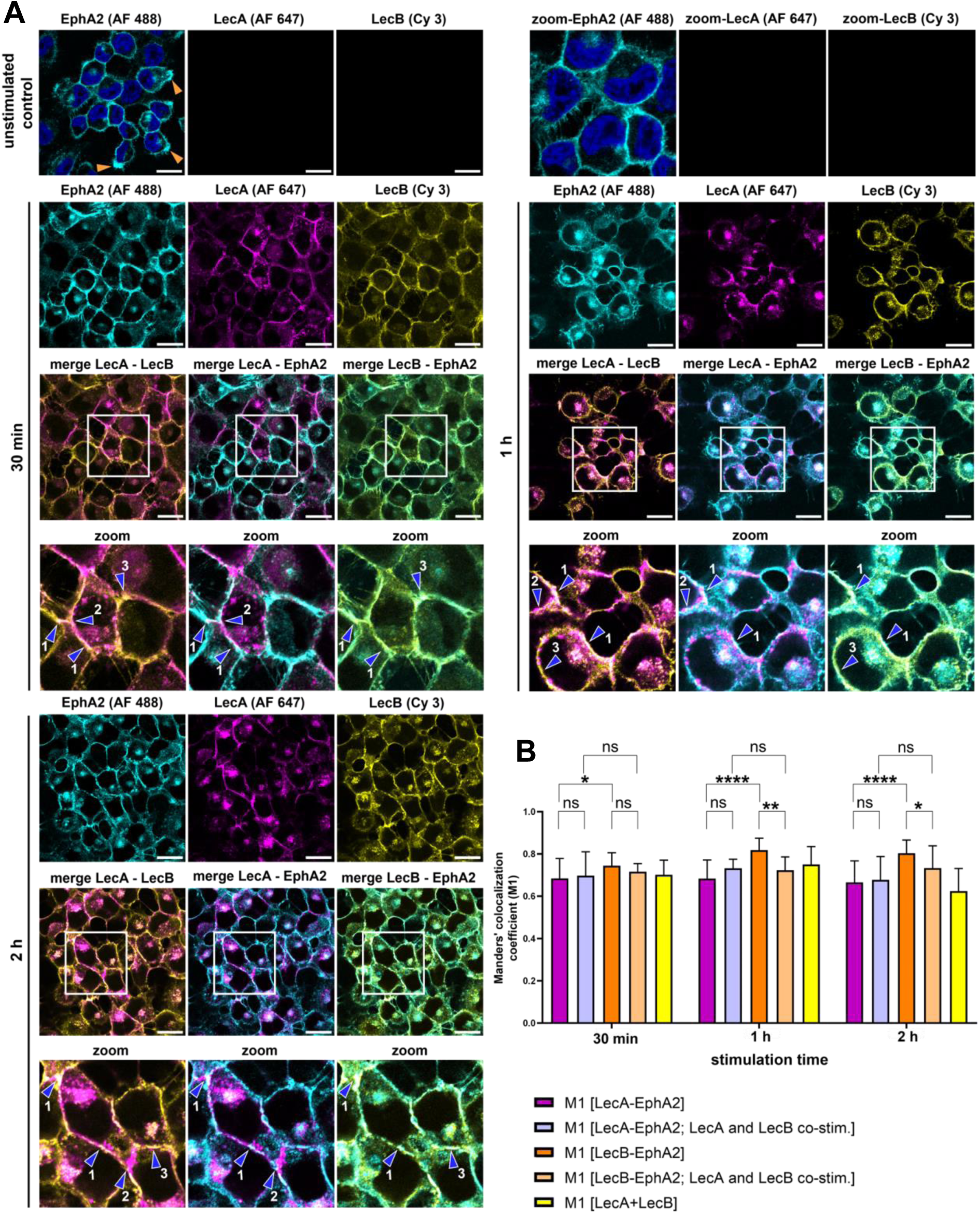
LecA and LecB colocalize with EphA2 at plasma membranes of H1299 cells. **(A)** Representative fluorescence microscopy images of H1299 cells co-stimulated with equimolar concentrations of LecA-AF 647 (pseudocolored in magenta) and LecB-Cy 3 (pseudocolored in yellow) for 30 min to 2 h at 37 °C, followed by fixation with 4% PFA and immunostaining for EphA2 (AF 488, pseudocolored in cyan). Nuclei were counterstained with DAPI (blue). In unstimulated control cells, orange arrowheads indicate EphA2 clusters at lamellipodia. Numbered dark blue arrowheads highlight colocalized signals (white overlay in merged images) between LecA, LecB, and EphA2 (1), LecA and EphA2 only (2), and LecB and EphA2 only (3). Scale bars, 20 µm. **(B)** Quantification of Manders’ colocalization coefficient 1 (M1). LecA-EphA2 and LecB-EphA2 indicate M1 values from single-lectin stimulations; LecA-EphA2 (co-stim.) and LecB-EphA2 (co-stim.) indicate M1 values from co-stimulation experiments; LecA + LecB indicates the M1 value between LecA and LecB in co-stimulation experiments. Data are presented as mean ± SD of 8–35 cells per condition from two independent experiments (*n* = 2). Statistical significance was determined by ordinary two-way ANOVA followed by Šídák’s multiple comparisons test (ns, not significant; *p < 0.05; **p < 0.01; ***p < 0.001; ****p < 0.0001).

In unstimulated cells, the EphA2 signal was distributed fairly evenly along the plasma membrane, showing prominent foci at lamellipodia (Fig. 3A and Supplementary Fig. 2, unstimulated control, EphA2-AF488, cyan, orange arrowheads). Across all conditions, distinct intracellular EphA2-AF488 clusters were also visible in the perinuclear area, likely representing receptor transit through the ER–Golgi secretory pathway. Upon single-lectin stimulation, both lectins strongly correlated with EphA2 at the plasma membrane over the entire stimulation period, with LecB displaying greater colocalization than LecA (Fig. 3B, LecA-EphA2/LecB-EphA2, and Supplementary Fig. 2). Accordingly, M1 MCC values reached 0.70 for LecA-EphA2 and 0.75-0.80 for LecB-EphA2, while M2 values reached 0.60 for EphA2-LecA and 0.80 for EphA2-LecB (Supplementary Fig. 3). The comparable M1 and M2 values across conditions indicate similar plasma membrane abundances of the lectins and EphA2, demonstrating that both lectins bind to targets within the same EphA2-enriched membrane domains.

Co-stimulation of H1299 cells with LecA and LecB maintained the spatial association of each lectin with EphA2 at the plasma membrane, yielding M1 MCCs of 0.65-0.75 for LecA and 0.70-0.80 for LecB (Fig. 3B, LecA-EphA2 and LecB-EphA2; LecA and LecB co-stim.). This demonstrates that neither lectin displaces the other from EphA2 membrane domains, ruling out apparent competitive binding for their plasma membrane targets. While distinct regions showed exclusive overlap between EphA2-AF488 and either LecA-AF647 (Fig. 3A, No. 2, dark blue arrowheads) or LecB-Cy3 (Fig. 3A, No. 3, dark blue arrowheads), LecA, LecB, and EphA2 predominantly overlaid within the same membrane domains (Fig. 3A, No. 1, dark blue arrowheads). Furthermore, LecA and LecB strongly colocalized with one another (Fig. 3B, M1 MCCs of 0.65-0.75; LecA + LecB), supporting the notion that both lectins target the same membrane domains and potentially act in concert at the EphA2 receptor.

Intriguingly, we also observed regions of mutually exclusive distribution at the plasma membrane, most notably for LecA and EphA2 compared with LecB and EphA2 (Fig. 3A and Supplementary Fig. 2, orange arrowheads). This suggests that lectin binding can actively remodel the spatial organization of EphA2 at the plasma membrane, potentially inducing receptor clustering or segregation to modulate downstream signaling.

### LecA and LecB interact differently with the EphA2 receptor

Since LecA and LecB exhibit a high degree of colocalization with EphA2 at the plasma membrane, we next examined whether EphA2 belongs to the host cell surface interactome targeted by these lectins. In particular, we assessed whether either lectin interacts directly or indirectly with EphA2 by combining pull-down assays, immunoprecipitation (IP), lectin blots, and protein docking simulations followed by molecular dynamics (MD) equilibration.

Pull-down assays identified EphA2 as a potential interaction partner of LecB, with both proteins co-eluting in the extract fraction (Fig. 4A, extract). Comparable amounts of EphA2 and LecB were recovered at 1 h and 3 h, likely because shifting cells from ice to 37 °C without medium replacement enabled remaining unbound lectins to bind the plasma membrane continuously. Meanwhile, the presence of both proteins in the supernatant indicates that only a fraction of EphA2 belongs to the LecB interactome and that not all LecB localizes to EphA2 membrane domains (Fig. 4A, supernatant). Conversely, pull-downs of the LecA-targeted membrane interactome yielded markedly less EphA2 than those with LecB (Supplementary Fig. 4, extract). Although the absence of LecA in the supernatant confirms efficient capture of LecA complexes, residual EphA2 in this fraction, analogous to LecB, suggests that only a minor portion of EphA2 associates with the LecA interactome. Moreover, detection of EphA2 in control extracts reveals non-specific background binding to streptavidin beads, further indicating a weak or transient interaction with LecA.

**Fig. 4.**
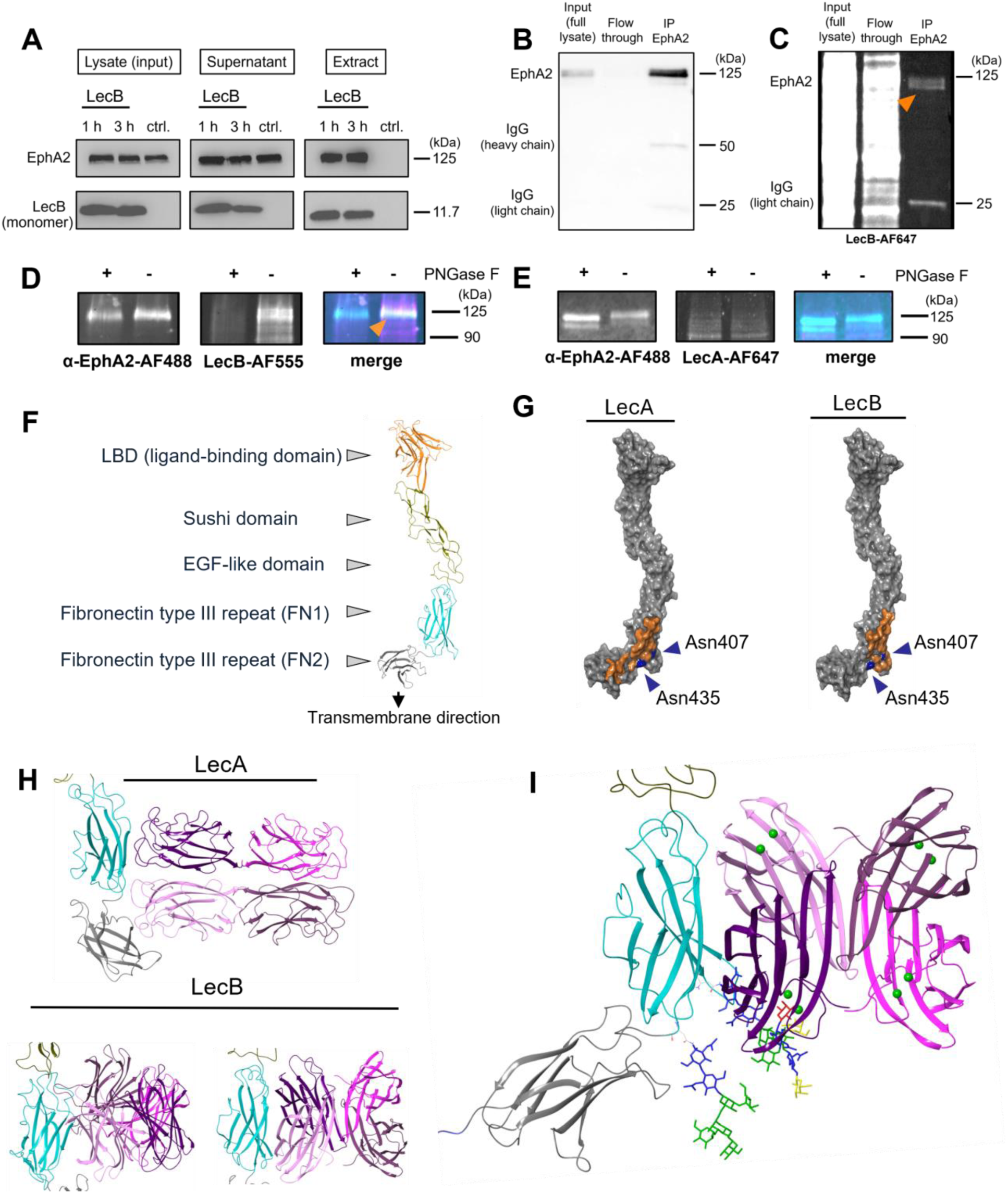
Lectins exploit protein–carbohydrate and protein–protein interactions to a different extent upon EphA2 engagement. **(A)** Representative Western blot images from pull-down assays of LecB and EphA2. H1299 cells were incubated with biotinylated LecB for 1 h or 3 h at 37 °C, whereas control cells remained untreated. Lysate, whole-cell lysate prior to incubation with magnetic streptavidin-coated beads; supernatant, unbound fraction collected after bead incubation; extract, eluted LecB together with its direct or indirect interaction partners (here, EphA2). **(B)** Representative Western blot image of immuno-precipitated EphA2 (IP-EphA2). **(C)** Lectin blot of IP-EphA2 samples probed with LecB-AF647. Co-detection of LecB and EphA2 signals is indicated by the orange arrowhead. Detection of IgG heavy and light chains results from secondary antibody binding to the precipitating IgG. **(D, E)** Representative lectin blots of cell lysates with (+) or without (–) PNGase F treatment. Membranes were first incubated with LecB-AF555 (D) or LecA-AF647 (E), followed by an anti-EphA2 primary antibody and an AF488-conjugated secondary antibody. Merged signals of magenta (LecB or LecA) and cyan (EphA2) appear as a white band (D, merge, orange arrowhead). **(F, G)** Structure of the EphA2 ectodomain illustrated as a ribbon model (F) and as grey surface representation (G). Individual domains are color-coded, except for the Sushi and EGF-like domains; the black arrow indicates the orientation toward the transmembrane region (F). Potential binding epitopes for LecA and LecB are highlighted in orange on the FN1 and FN2 domains, with Asn407 and Asn435 shown in blue (G). **(H)** Ribbon models illustrating protein–protein docking poses of LecA (top) and LecB (bottom) with the EphA2 FN1 and/or FN2 domains. Lectin monomers are colored in shades of magenta, while EphA2 domains follow the color scheme in (F). **(I)** Ribbon model of the LecB–EphA2 complex stabilized through both protein–carbohydrate and protein–protein interactions. Glycan structures are colored according to the Symbol Nomenclature for Glycans (SNFG), and calcium ions are depicted as green spheres.

To investigate whether LecA or LecB directly binds to EphA2 in a carbohydrate-dependent manner, we next performed IP experiments and lectin blots. Furthermore, samples were treated with or without PNGase F to evaluate how N-glycan removal affects lectin binding to EphA2.

Immunoprecipitation of whole-cell lysates from unstimulated cells yielded a distinct EphA2 band in the IP fraction (Fig. 4B, IP EphA2). A control IP targeting β1-integrin, which shares a similar molecular weight with EphA2 (∼125-130 kDa), showed no detectable EphA2 signal (Supplementary Fig. 5), confirming the specificity of the anti-EphA2 antibody and the IP protocol. The IP samples were then probed in lectin blots with fluorescently labeled LecA-AF647 or LecB-AF647, alongside immunostaining for EphA2 using an anti-EphA2 primary antibody and an AF488-conjugated secondary antibody. Analysis of the merged fluorescence signals revealed a distinct overlap between LecB and EphA2, demonstrating a direct carbohydrate-dependent interaction (Fig. 4C, orange arrowhead), whereas no signal was observed for LecA (Supplementary Fig. 6).

PNGase F treatment reduced the apparent molecular weight of EphA2 by ∼5–10 kDa (Fig. 4D, E, anti-EphA2/AF488), consistent with the removal of N-linked glycans from its two known N-glycosylation sites at Asn407 (FN1 domain) and Asn435 (FN2 domain) (25,57). While untreated lysates showed strong LecB binding to multiple glycosylated proteins, LecB reactivity was nearly abolished upon PNGase F treatment (Fig. 4D, LecB-AF555). Crucially, untreated samples exhibited a distinct signal overlap between LecB-AF555 and EphA2-AF488 (Fig. 4D, merge, orange arrowhead), confirming that LecB binds N-glycans on a ∼125 kDa protein corresponding to EphA2. Conversely, no overlap was observed between LecA-AF647 and anti-EphA2/AF488 (Fig. 4E, merge). Combined with our IP data, these results indicate that LecA does not directly engage the glycans of EphA2.

Potential carbohydrate-dependent and-independent interaction modes of the lectins with EphA2 were further explored using protein–protein docking simulations followed by MD equilibration. Initial docking searches for potential interaction epitopes revealed a preference of both lectins for the FN2 domain and the LBD of EphA2 (Fig. 4F). However, subsequent equilibration of the docked complexes over 400 ns MD trajectories in explicit water (Supplementary Table 1 and Supplementary Fig. 7; RMSD, interacting residues and binding energies) confirmed the stability of only one complex for LecA and two complexes for LecB, with interactions occurring primarily at the FN1 domain in all cases (Fig. 4G, orange surfaces and Fig. 4H).

Surface representations in Fig. 4G (blue areas) highlight potential lectin-binding glycosylation sites within the FN1 and FN2 domains of EphA2. These domains have been reported to carry diverse glycan structures, including fucosylated complex glycans (58) in the FN1 domain and oligomannose-type glycans in the FN2 domain (Fig. 4I). To evaluate the influence of N-glycans on lectin binding, we modeled a glycosylated variant of EphA2 carrying a terminal fucose residue on an antennary branch attached to Asn407. This modification enabled direct engagement by LecB (Fig. 4I, FN1), consistent with our experimentally confirmed protein–carbohydrate interaction between LecB and EphA2. Notably, the energetically most favorable EphA2–LecB docking complex maintains the optimal spatial distance between the N-glycan and the fucose-binding pocket of LecB, allowing the terminal fucose to adopt a binding orientation consistent with experimentally determined glycan-bound crystal structures (PDB code: 5A3O) (59,60).

Taken together, LecB can directly bind EphA2, most likely by recognizing fucosylated epitopes on complex N-glycans within the receptor’s FN1 domain. Conversely, LecA does not directly bind EphA2 glycans, but may associate with the receptor’s FN1 and FN2 domains via weak protein– protein interactions.

### *P. aeruginosa* adhesion triggers local EphA2 clustering and induces kinase-dependent activation

Given that soluble LecA and LecB strongly colocalize with EphA2 at the host plasma membrane, with LecB directly engaging the receptor, we next investigated whether similar interactions occur during infection with live *P. aeruginosa*. Specifically, we investigated whether bacterial adhesion actively recruits EphA2 or preferentially targets pre-existing EphA2-enriched membrane domains. Furthermore, we assessed if live *P. aeruginosa* triggers EphA2 activation in a manner comparable to the soluble lectins. To address these questions, H1299 cells were challenged with fluorescently labeled *P. aeruginosa* (PAO1-mCherry), followed by EphA2 immunostaining (EphA2/AF488, cyan) and Western blot analysis of EphA2 Tyr588 phosphorylation.

Immunofluorescence imaging revealed distinct EphA2 clusters at *P. aeruginosa* adhesion sites, indicating a strong spatial association between the receptor and bacteria at the plasma membrane (Fig. 5A, zoom 1–3, white merge, orange arrowheads). Conversely, unstimulated control cells exhibited a fairly uniform EphA2/AF488 signal distribution across the plasma membrane, with virtually no detectable EphA2 clusters (Fig. 5A, unstimulated control, orange arrowheads).

**Fig. 5.**
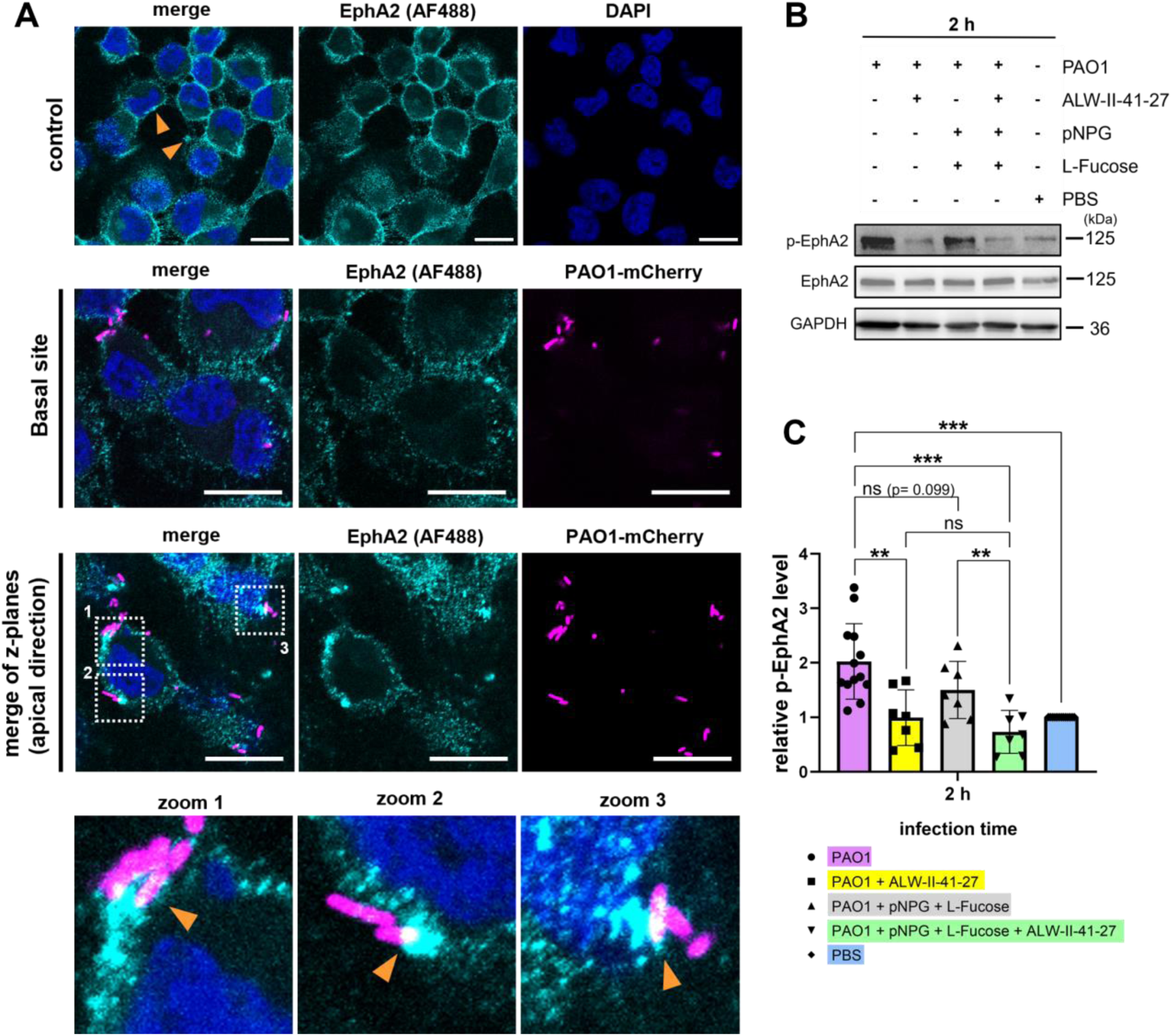
***P. aeruginosa* actively recruits EphA2 at the plasma membrane and induces kinase-dependent receptor activation. (A)** Representative confocal microscopy images of H1299 cells challenged with fluorescently labeled *P. aeruginosa* PAO1-mCherry (pseudocolored in magenta; MOI 15) for 2 h at 37 °C. Cells were subsequently fixed with 16% PFA, permeabilized, and stained with an anti-EphA2 primary antibody followed by an AF488-conjugated secondary antibody (pseudocolored in cyan). Magnified regions show plasma membrane areas with overlapping EphA2/AF488 and PAO1-mCherry signals (white merge, orange arrowheads), whereas untreated control cells displayed almost no EphA2 clustering (orange arrowheads). Scale bars, 20 µm. **(B)** Representative Western blot image showing pronounced p-EphA2 induction in H1299 cells challenged with *P. aeruginosa* (PAO1; MOI 15) for 2 h at 37 °C, and a strong reduction upon co-treatment with ALW-II-41-27. Cells were infected with bacteria alone or pre-treated with ALW-II-41-27 (500 nM, 45 min, 37 °C), followed by bacterial stimulation with freshly added inhibitor (500 nM). Where indicated, bacteria were pre-treated with pNPG (10 mM) and L-fucose (45 mM) for 30 min at RT prior to infection of untreated or ALW-II-41-27-pre-treated cells. PBS served as a vehicle control and GAPDH as a loading control. **(C)** Quantification and statistical analysis of p-EphA2 levels from independent Western blot experiments, including the representative blot shown in (B). Densitometric values were normalized to total protein load via Ponceau S staining and expressed relative to the PBS control. Data are presented as mean ± SD of *n* ≥ 7 biological replicates per group. Statistical analysis was performed using unpaired two-tailed t-tests (ns, not significant; *p < 0.05; **p < 0.01; ***p < 0.001; ****p < 0.0001).

Stimulation with PAO1 resulted in a significant increase in EphA2 phosphorylation compared with unstimulated control cells (p = 0.0002), demonstrating that live bacteria can indeed activate the receptor (Fig. 5B, C). This phosphorylation depended strongly on the receptor’s kinase activity, as simultaneous treatment with ALW-II-41-27 drastically attenuated p-EphA2 levels back to baseline PBS control levels (Fig. 5B, C; 100% reduction). Additionally, co-treatment with pNPG and L-fucose reduced PAO1-induced p-EphA2 levels by approximately 50% (Fig. 5C), aligning with previous studies showing that these competitive inhibitors disrupt lectin–host receptor interactions and thereby reduce bacterial invasiveness (15,21,55,61). The most profound reduction in p-EphA2 was observed upon simultaneous application of PAO1, pNPG, L-fucose, and ALW-II-41-27, with levels declining even below baseline control conditions (Fig. 5B, C). Collectively, these findings demonstrate that *P. aeruginosa* engages host glycans via its lectins to trigger kinase-dependent canonical EphA2 activation.

## Discussion

Bacterial pathogens have evolved sophisticated strategies to manipulate host cell signaling pathways, frequently exploiting host defense and regulatory mechanisms to facilitate cellular invasion, evade immune responses, and promote intracellular survival and replication (62–64). A recurring theme in host-pathogen interactions is the targeting of the host plasma membrane, where the strategic engagement of RTKs triggers the actin rearrangements necessary for bacterial entry. While the role of *P. aeruginosa* lectins LecA and LecB in binding to host cell surface glycans is well-documented (16,21), the mechanism by which this physical tethering is converted into an intracellular signal has remained a critical gap in our understanding.

In the present study, we demonstrate that soluble LecA and LecB strongly activate canonical EphA2 signaling via Tyr588 phosphorylation. This finding is particularly significant as it expands the known repertoire of RTKs leveraged by *P. aeruginosa.* Whereas previous studies focused on the activation of the platelet-derived growth factor receptor (PDGFR) and the epidermal growth factor receptor (EGFR) through pili or flagella (65,66), our results reveal a more diversified strategy for host cell manipulation. By engaging EphA2, *P. aeruginosa* lectins may co-opt signaling pathways distinct from those triggered by canonical pathogen-associated molecular patterns (PAMPs), potentially enabling the pathogen to subvert certain host immune defenses.

The strict dependence of EphA2 activation on lectin-carbohydrate binding, evidenced by the inhibitory effects of pNPG and L-fucose, underscores glycan recognition as a crucial initial step in triggering host signaling. This mirrors the behavior of other “lectin-like” adhesin-producing pathogens, such as *Vibrio cholerae* or *Helicobacter pylori*, which utilize specific carbohydrate signatures to anchor themselves to host plasma membranes prior to initiating downstream signaling cascades (67,68). Although LecA and LecB rely on carbohydrate binding to activate EphA2, their dependence on specific membrane components differs markedly. Glycosphingolipids are frequently exploited by bacterial pathogens as host receptors to facilitate adhesion, invasion, and toxin uptake (69,70). While GSL depletion significantly impaired LecA-induced EphA2 activation, LecB-induced activation remained largely unaffected. This suggests a functional specialization in which LecA promotes bacterial uptake primarily through Gb3 binding, whereas LecB activates EphA2 through a mechanism that relies less on GSLs.

The lectin-induced downstream phosphorylation of CrkII at Tyr221 further links lectin binding to the host machinery facilitating bacterial invasion. Modulation of Crk/CrkII adapter proteins is a hallmark of invasion strategies employed by a wide array of pathogens, including *Yersinia pseudotuberculosis*, *Shigella flexneri*, *Streptococcus agalactiae*, and *P. aeruginosa* (16,71–74). Interestingly, our data reveal a mechanistic dichotomy within *P. aeruginosa* itself: whereas the type III secretion effector ExoT suppresses CrkII to dampen inflammatory responses (75–77), LecB predominantly promotes CrkII phosphorylation. Treatment with the EphA2 inhibitor ALW-II-41-27 confirmed that this process is kinase-dependent, aligning it with the mechanism observed during Kaposi’s sarcoma-associated herpesvirus (KSHV) infection. Specifically, KSHV activates p130Cas via EphA2, leading to the association of p130Cas and Crk with phosphorylated EphA2 (78). Thus, our findings further establish EphA2 as a pivotal mediator of host-pathogen interactions.

One of the most intriguing aspects of our findings is how these lectins trigger EphA2 activation in the absence of its canonical ligand, ephrin-A1. Because canonical EphA2 signaling depends heavily on receptor clustering and the formation of higher-order complexes (79,80), the colocalization and potentiating effects of both LecA and LecB point to a mechanism driven by their multivalent nature. We propose that LecA and LecB bind to glycosylated plasma membrane targets, such as Gb3 or integrins, in close proximity to EphA2, thereby altering the receptor’s lateral membrane distribution. Since both lectins can cross-link membrane receptors via their opposing binding pockets (18,81,82), and given our previous findings that LecA can disperse liquid-ordered domains (83), we hypothesize that lectin-triggered membrane reorganization modulates EphA2 clustering or lateral dispersion. This remodeling may promote the assembly of biomolecular condensates or specialized membrane-associated signaling platforms through receptor clustering and phase separation (84,25,85). Similar organizational principles have been described in other cellular contexts, where increased local receptor density, membrane confinement, or partitioning into lipid nanodomains drives ligand-independent or constitutive signaling (86–88). In this sense, *P. aeruginosa* does not merely adhere to the host cell; rather, it actively remodels the plasma membrane topography to mimic the signaling microenvironment generated by ephrin-Eph interactions.

Our mechanistic investigations further clarify the distinct interaction modes between these lectins and EphA2. The direct binding of LecB to N-linked glycans on EphA2, contrasted with the weak protein–protein interactions likely mediated by the receptor’s FN1 and FN2 domains for LecA, suggests a complementary or cooperative effect on the host cell. This functional interplay endows the pathogen with considerable versatility; whereas LecA exploits its high affinity for Gb3 to drive membrane engulfment (15), LecB engages a broader repertoire of glycoconjugates (89–92). Such functional redundancy in bacterial adhesion provides *P. aeruginosa* with alternative host-recognition strategies, potentially facilitating colonization across tissues with diverse glycosylation profiles (93).

Finally, translating these soluble lectin findings to live *P. aeruginosa* infection establishes a compelling link to bacterial pathogenesis. The clustering of EphA2 at the bacterium-host interface indicates that the living pathogen actively engages this receptor during adhesion. This conclusion is reinforced by our observation that *P. aeruginosa* infection significantly induces EphA2 activation in a kinase-and partially carbohydrate-dependent manner, identifying LecA and LecB as key drivers of EphA2 signaling during host-pathogen association. Notably, our findings complement the previously described role of flagellin, which triggers TLR5-dependent ephrin-A1 cleavage via ADAM9 to induce ligand-independent EphA2 signaling in airway epithelial cells (94).

Collectively, these observations reinforce the emerging paradigm that EphA2 serves as a conserved entry and signaling receptor for diverse pathogens including *Chlamydia trachomatis*, *Helicobacter pylori*, uropathogenic *Escherichia coli*, *Klebsiella pneumoniae*, and *Enterococcus faecalis* (33,37,95). However, the specific bacterial determinants responsible for triggering EphA2 activation have remained largely undefined, with the notable exception of *C. trachomatis*, whose elementary bodies directly engage the receptor to promote co-internalization of the receptor-pathogen complex (37). By identifying LecA-and LecB-driven EphA2 activation, our work highlights the lectin-EphA2 axis as a promising target for anti-virulence intervention. Disrupting this axis, either via glycomimetic lectin inhibitors or targeted kinase blockage, represents an attractive therapeutic strategy to attenuate *P. aeruginosa* infection.

## Conclusion

In summary, our study uncovers a previously unrecognized signaling axis in which the *Pseudomonas aeruginosa* lectins LecA and LecB hijack the host RTK EphA2 to promote bacterial invasion. Both lectins trigger canonical EphA2 signaling through Tyr588 phosphorylation and downstream CrkII activation. Although LecA elicits stronger EphA2 phosphorylation than LecB at equimolar concentrations, co-stimulation with both lectins further amplifies receptor activation, indicating a cooperative and additive effect. We therefore propose distinct yet complementary modes of EphA2 engagement: LecA primarily activates EphA2 through carbohydrate-dependent interactions with Gb3 combined with weaker LecA–EphA2 protein–protein binding, whereas LecB binds directly to N-linked glycans on EphA2. Both interaction modes drive lateral membrane reorganization that alters the stability of EphA2-containing membrane domains, promoting conformational changes and subsequent receptor activation. Crucially, live *P. aeruginosa* exploits this lectin–EphA2 signaling platform at the bacterium–host interface during adhesion and entry. Together, these findings expand our understanding of how bacterial lectins translate extracellular glycan recognition into intracellular host signaling and highlight the lectin–EphA2 axis as a promising target for anti-virulence therapies against *P. aeruginosa*.

## Abbreviations

AF: Alexa Fluor
ANOVA: Analysis of variance
BSA: Bovine serum albumin
BCA: Bicinchoninic acid
CrkII: CT10 regulator of kinase II
Cy3: Cyanine 3
DABCO: 1,4-diazabicyclo [2.2.2] octane
EGFR: Epidermal growth factor receptor
Eph: Erythropoietin-producing hepatocellular
F-actin: Filamentous actin
FCS: Fetal calf serum
FN1: Fibronectin type III domain 1
FN2: Fibronectin type III domain 2
Gb3: Globotriaosylceramide, CD77, P^k^ blood group antigen
GEF: Guanine nucleotide exchange factor
GlcCer: Glucosylceramide
GPI: Glycosylphosphatidylinositol
GSL: Glycosphingolipid
HABA: 4′-hydroxyazobenzene-2-carboxylic acid
H1299: Human non-small cell lung carcinoma cell line
IP: Immunoprecipitation
JM: Juxtamembrane
KSHV: Kaposi’s sarcoma-associated herpesvirus
LB: Luria-Bertani
LBD: Ligand binding domain
LINCS: LINear Constraint Solver
MCC: Manders’ colocalization coefficient
MD: Molecular dynamics
MDCK: Madin-Darby canine kidney cells
MM/GBSA: Molecular mechanics/generalized born surface area
MOI: Multiplicity of infection
MW: Molecular weight
NHS: N-hydroxysuccinimide
PAMP: Pathogen-associated molecular pattern
PAO1: P. aeruginosa wild-type strain
PDGFR: Platelet-derived growth factor receptor
PDZ: PSD95/Dlg1/ZO-1 domain
PIP₃: Phosphatidylinositol (3,4,5)-trisphosphate
pNPG: p-nitrophenyl α-D-galactopyranoside
PME: Particle Mesh Ewald
PNGase F: Peptide:N-glycosidase F
PPMP: D-threo-1-phenyl-2-palmitoylamino-3-morpholino-1-propanol
RIPA: Radio-immunoprecipitation assay
RMSD: Root-mean-square deviation
ROI: Region of interest
RT: Room temperature
RTK: Receptor tyrosine kinase
SAM: Sterile alpha motif
SD: Standard deviation
StxB: Shiga toxin B-subunit
TM: Transmembrane domain
WT: Wild-type

## Declarations

### Data availability

The datasets supporting the conclusions of this article are available in the https://freidata.uni-freiburg.de repository, DOI: 10.60493/5v4b7-mq882.

## Acknowledgements

The authors acknowledge support by the state of Baden-Württemberg through bwHPC and the German Research Foundation (DFG) through grant INST 35/1597-1 FUGG. We would like to particularly thank Dr. Anne Imberty (CNRS, Grenoble, France) for kindly providing the plasmids pET25-*pa1l* for LecA and pET25-*pa2l* for LecB production, the group of Prof. Dr. Karl-Erich Jäger (HHU Düsseldorf, Germany) for kindly providing the PAO1 wt strain and the group of Prof. Dr. Rolf Kümmerli (Department of Quantitative Biomedicine, Zurich, Switzerland) for kindly providing a constitutively expressing PAO1-mCherry strain for our immunofluorescence experiments. We also gratefully acknowledge Lena Wolter for excellent assistance in daily lab routine and Jonas Gräber for experimental assistance. CS thanks the Spemann Graduate School of Biology and Medicine for great support.

## Research funding

This study was supported by the German Research Foundation (DFG) under Germany’s Excellence Strategy (CIBSS – EXC-2189 – Project ID 390939984) and the DFG grant Major Research Instrumentation (438033605), as well as the Ministry of Science, Research and the Arts of the State of Baden-Württemberg (Az: 33–7532.20).

## Authors’ contributions

The concept and design of the research was conducted by C.S. and W.R. Funding was acquired by W.R. C.S. performed, analyzed and interpreted all Western blot data regarding EphA2 and CrkII phosphorylation, pull-down experiments with LecB stimulations, LecA and LecB lectin blots with PNGase F treatment, immunofluorescence imaging, bacterial stimulation experiments and bacterial imaging and all statistical analyses. O.M. and F.N. performed and analyzed protein docking and MD simulations for protein–protein and protein–carbohydrate interaction studies of LecA, LecB and EphA2. A.S. performed immunoprecipitation experiments and lectin blots for LecA, LecB and EphA2 interaction studies. F.R. and M.X. performed pre-studies for LecA and EphA2 colocalization and interaction analysis. M.X. performed and analyzed pull-down experiments with LecA and EphA2. P.R.M. assisted with scientific expertise during pull-down and IP experiments, and colocalization analysis. The original draft was written by C.S., review and editing were performed by all authors. All authors agreed on the final version of the manuscript.

## Declarations of Generative AI and AI-assisted technologies in the writing process

During the preparation of this manuscript, the authors used ChatGPT (OpenAI, GPT-5.6 Luna) for language editing and refinement. The authors reviewed and verified all AI-assisted content and take full responsibility for the final content of the manuscript.

