## Supplementary information for "*Pseudomonas aeruginosa* lectins LecA and LecB differentially engage and activate the EphA2 receptor"

^†^ Former member of 1

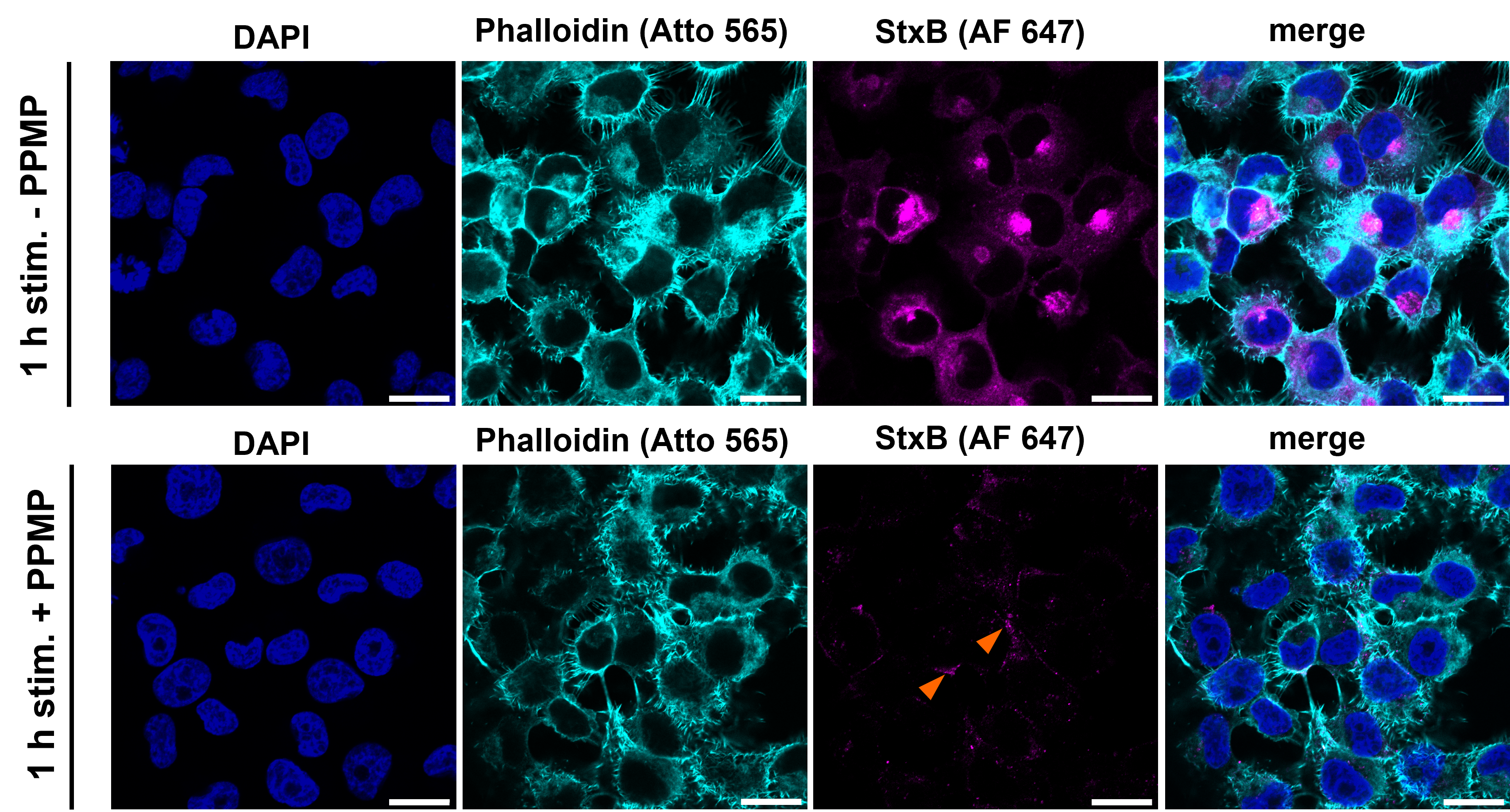

**A**

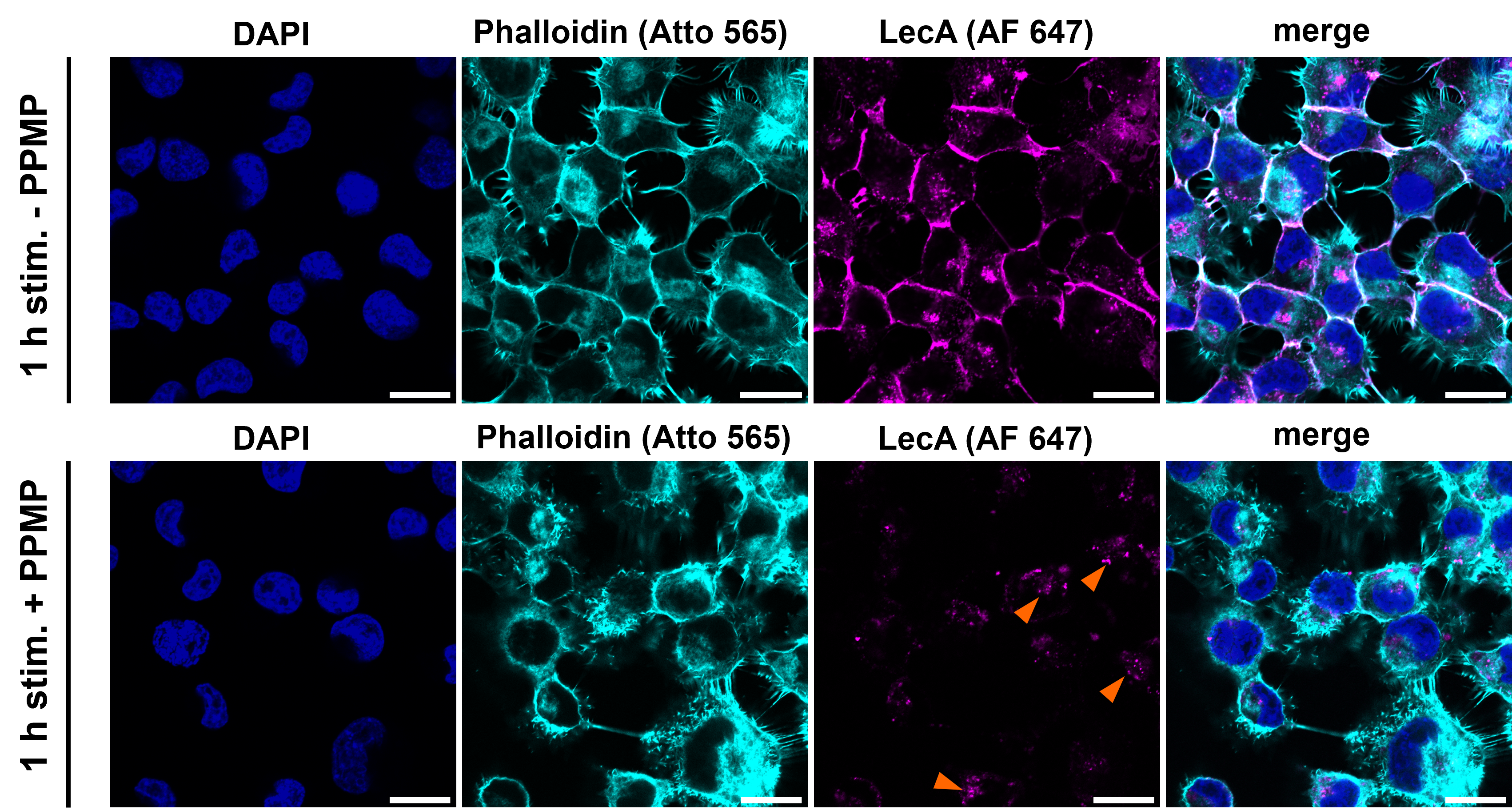

**B**

**Supplementary Fig. 1 Glycosphingolipid depletion strongly reduces plasma membrane binding of StxB and LecA to H1299 cells.**

**(A, B)** Representative immunofluorescence images of H1299 cells incubated for 1 h at 37 ℃ with either 260 nM of the Gb3-specific StxB (concentration based on the pentameric MW, AF 647, in magenta) (A) or 300 nM LecA (concentration based on the tetrameric MW, AF 647, in magenta) (B). Prior to stimulation, cells were treated with 2.5 µM PPMP (+) for 4 d or left untreated (–). The DNA was counterstained with DAPI (blue), and the actin cytoskeleton was stained with Phalloidin (Atto 565, in cyan) for spatial orientation. Merged images display all three fluorescence channels. Untreated cells exhibit strong binding and internalization of StxB and LecA in contrast to PPMP-treated cells. Orange arrowheads point to residual binding of the StxB (A) and LecA (B) to PPMP-treated cells. stim. = stimulation. Scale bars, 20 µm.

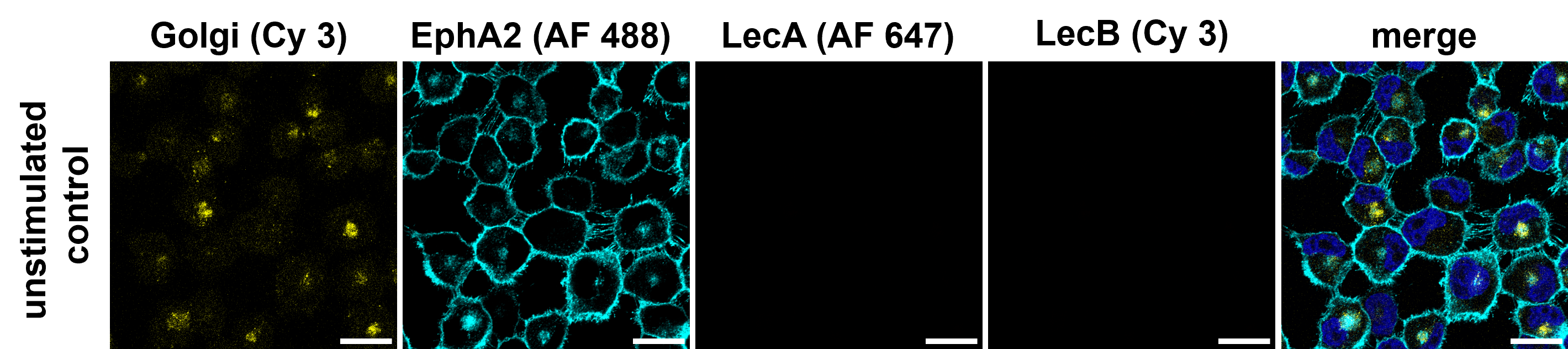

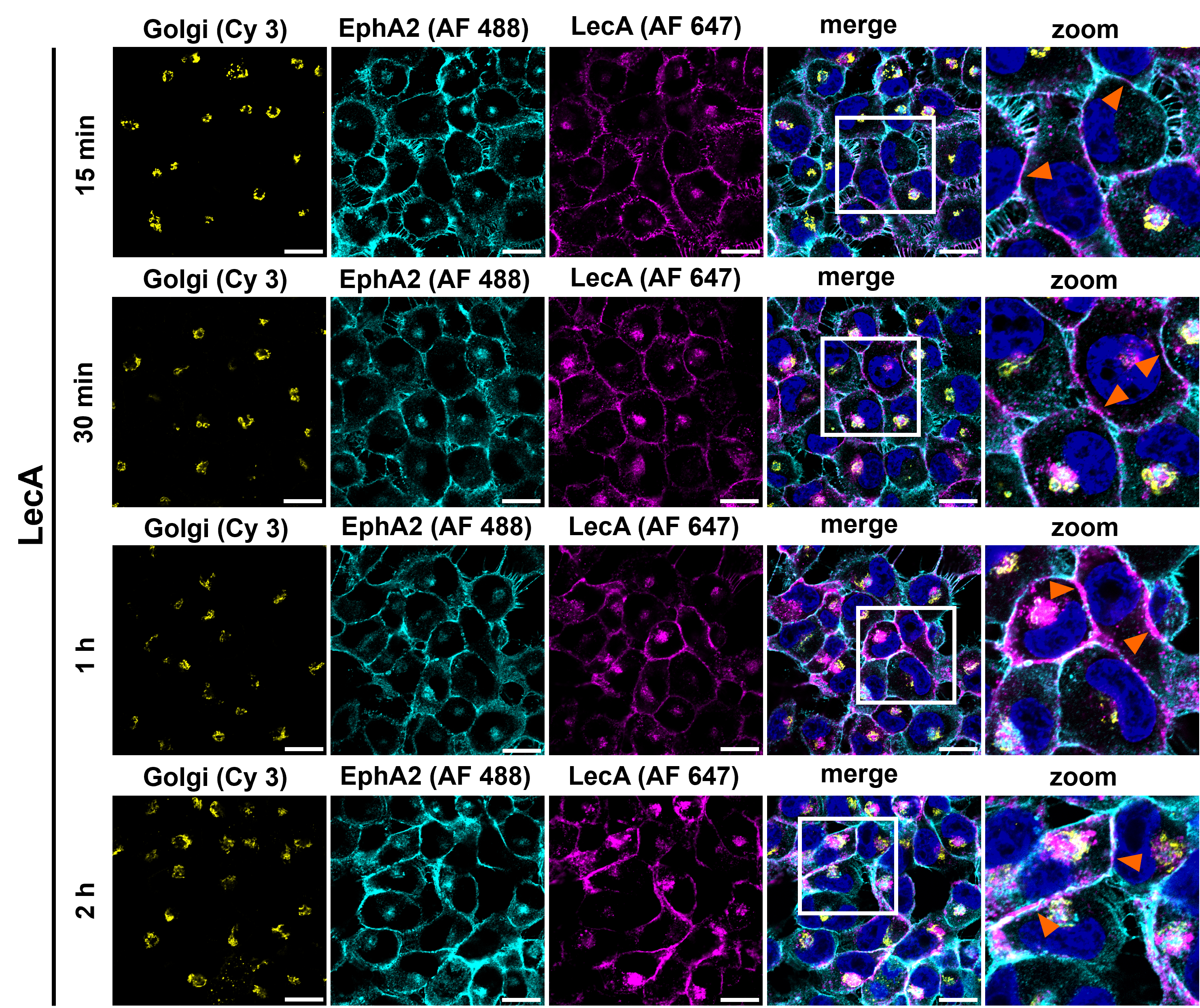

**A**

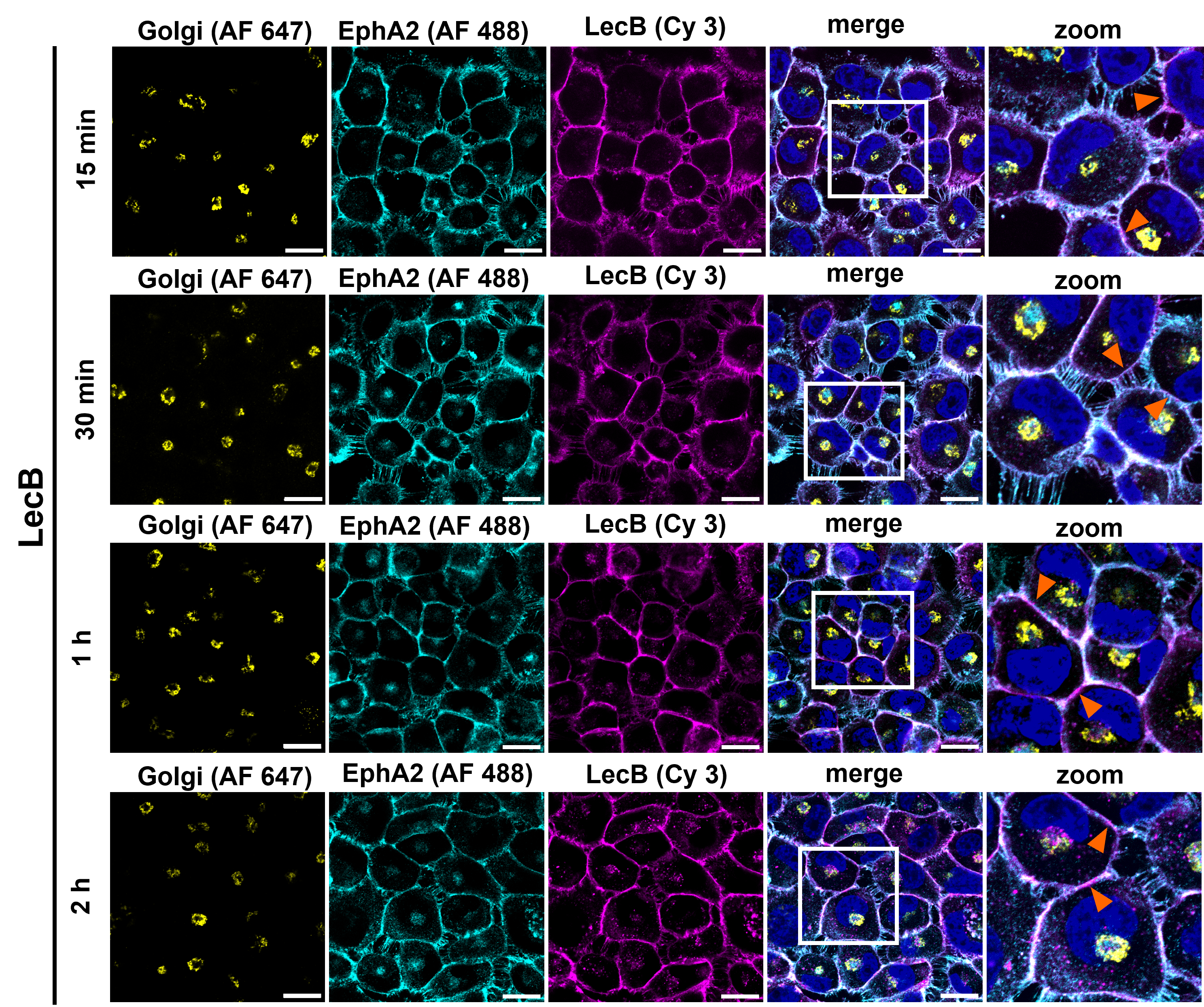

**B**

**Supplementary Fig. 2 LecA and LecB stimulation leads to a displacement of EphA2 at the plasma membrane.**

**(A, B)** Representative fluorescence images of lectin-stimulated H1299 cells stained for the cis/medial Golgi marker Giantin (Cy 3 or AF 647, in yellow, for intracellular spatial orientation) and EphA2 (AF 488, in cyan). The DNA was counterstained with DAPI (blue). Cells were stimulated at 37 ℃ over a time course of 15 min to 2 h with either 300 nM LecA-AF 647 (in magenta) (A) or 1 µM LecB-Cy 3 (in magenta) (B) (concentrations based on the tetrameric MW of each lectin), followed by fixation with 4% PFA. Control cells remained unstimulated. Merged images display all four fluorescence channels. In control cells, EphA2 remained evenly distributed along the plasma membrane. LecA stimulation resulted in a pronounced mutually exclusive (“either-or”) distribution of LecA and EphA2 signals at the plasma membrane (magnified insets, orange arrowheads) at early time points (15–30 min) and increasing overlay of these signals at later time points (2 h). In contrast, this mutually exclusive pattern was observed less frequently for LecB and EphA2 across all time points (magnified insets, orange arrowheads). Overall, LecB stimulation resulted in enhanced signal overlays of LecB and EphA2 compared to LecA and EphA2. Scale bars, 20 µm.

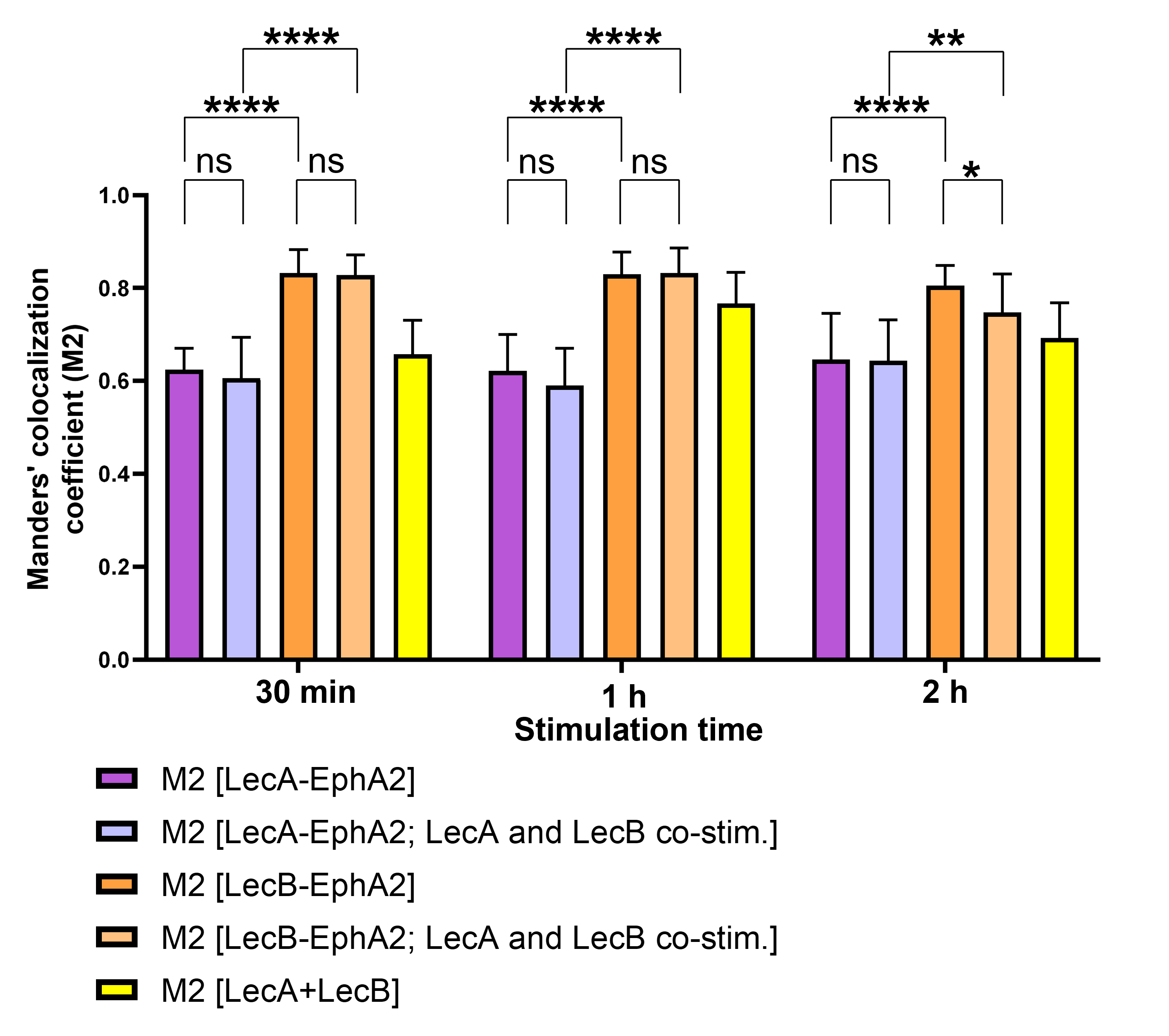

**Supplementary Fig. 3 Manders’ M2 coefficient reveals differential EphA2 colocalization with LecA and LecB during single- and co-stimulation.**

Quantification of Manders’ colocalization coefficient M2 from lectin stimulation experiments shown in Fig. 3. LecA-EphA2 and LecB-EphA2 represent Manders’ coefficients for EphA2 signal overlay with LecA or LecB, respectively, following single-lectin stimulation. LecA-EphA2 (co-stim.) and LecB-EphA2 (co-stim.) indicate Manders’ coefficients for EphA2 signal overlay with LecA or LecB, respectively in co-stimulation experiments. LecA + LecB represents the Manders’ coefficient between LecA and LecB upon co-stimulation. Statistical analysis was performed using an ordinary two-way ANOVA followed by Šídák’s multiple comparisons test. Data are presented as mean ± SD from *n* = 8–35 cells per condition across two independent experiments (ns, not significant; *p < 0.05, **p < 0.01, ***p < 0.001, ****p < 0.0001).

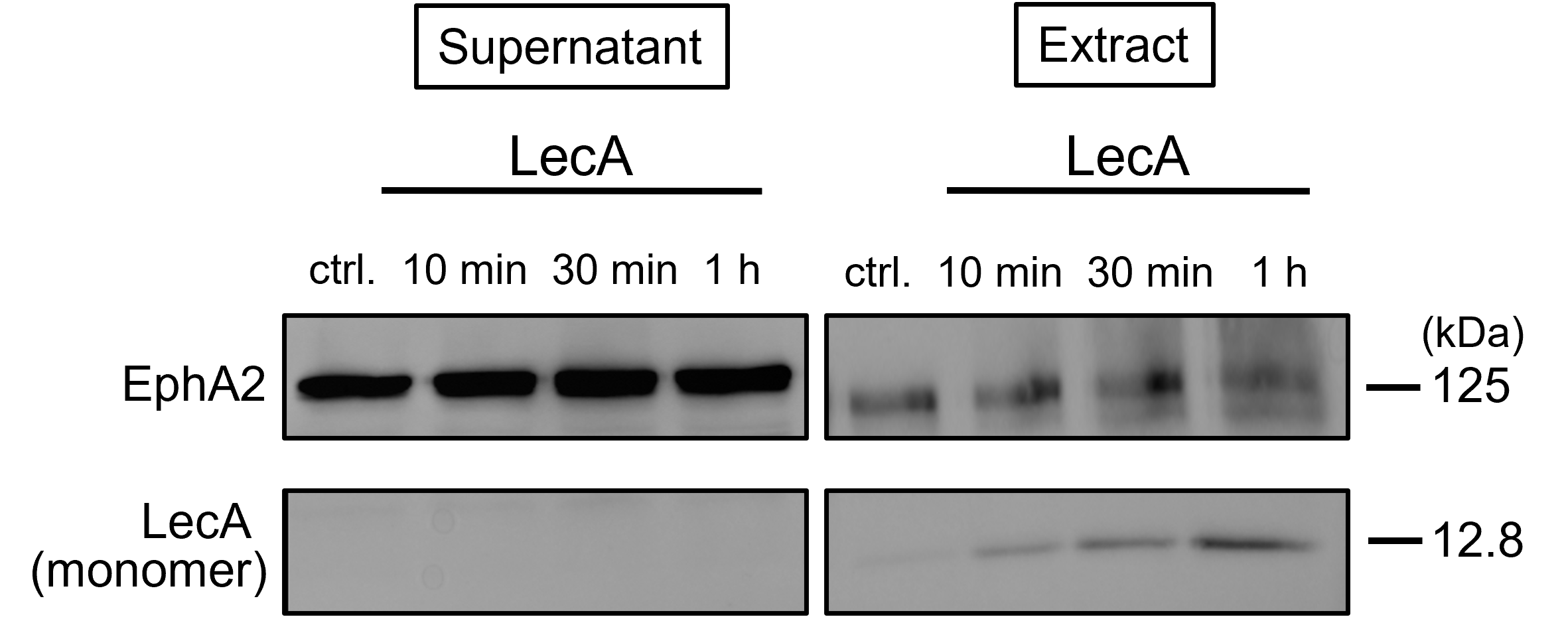

**Supplementary Fig. 4 EphA2 is weakly associated with the LecA membrane interactome**.

Representative immunoblots of streptavidin pull-down experiments analyzing the interaction between LecA and EphA2. H1299 cells were stimulated with 100 nM biotinylated LecA (LecA-biotin; concentration based on the tetrameric MW) for 10 min, 30 min, or 1 h at 37 ℃, whereas control cells remained unstimulated. Supernatant, unbound fraction collected after incubation with magnetic streptavidin-coated beads; extract, eluted fraction containing LecA-biotin and its direct or indirect interaction partners (here, EphA2). The absence of a LecA signal in the supernatant indicates efficient pull-down by the streptavidin-coated beads, whereas the strong residual EphA2 signal in the supernatant indicates that only a minor fraction of EphA2 associates with the LecA-targeted membrane interactome. The faint EphA2 band observed in the unstimulated control extract reflects nonspecific background binding to the beads.

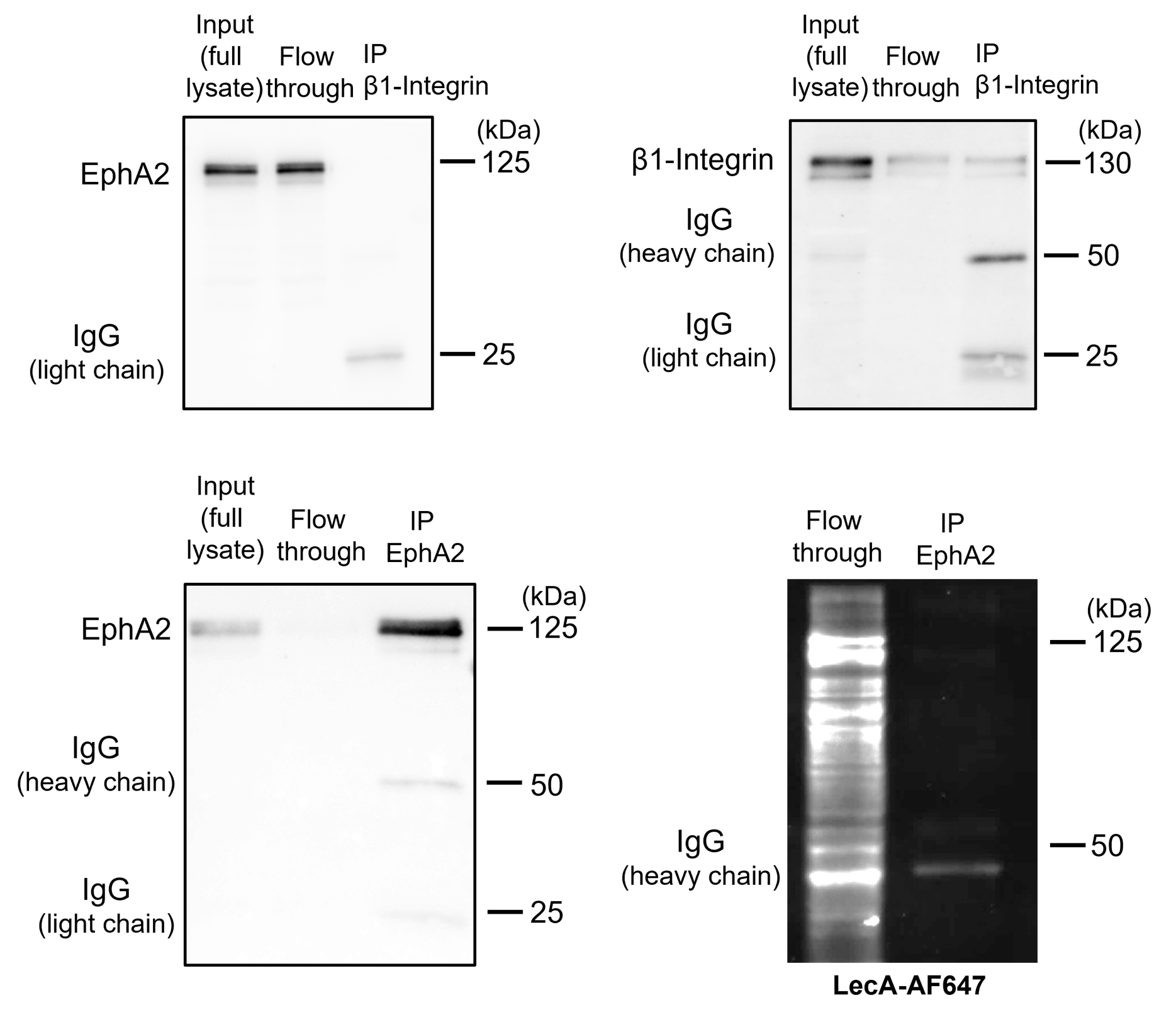

**Supplementary Fig. 5 EphA2-specific antibodies do not cross-react with immunoprecipitated β1-integrin.**

Whole-cell lysates of H1299 cells were incubated with β1-integrin–specific antibodies and immunoprecipitated using superparamagnetic Protein G Dynabeads. The whole-cell lysate (input), flow-through (unbound fraction prior to elution), and eluted antibody–antigen complexes (IP) were analyzed by immunoblotting using anti-EphA2 (left) and anti-β1-integrin (right) primary antibodies followed by HRP-conjugated secondary antibodies. Probing with anti-EphA2 antibodies yielded no detectable band in the β1-integrin IP fraction, whereas probing with anti-β1-integrin antibodies detected a prominent band in the corresponding IP fraction (right, orange arrowhead).

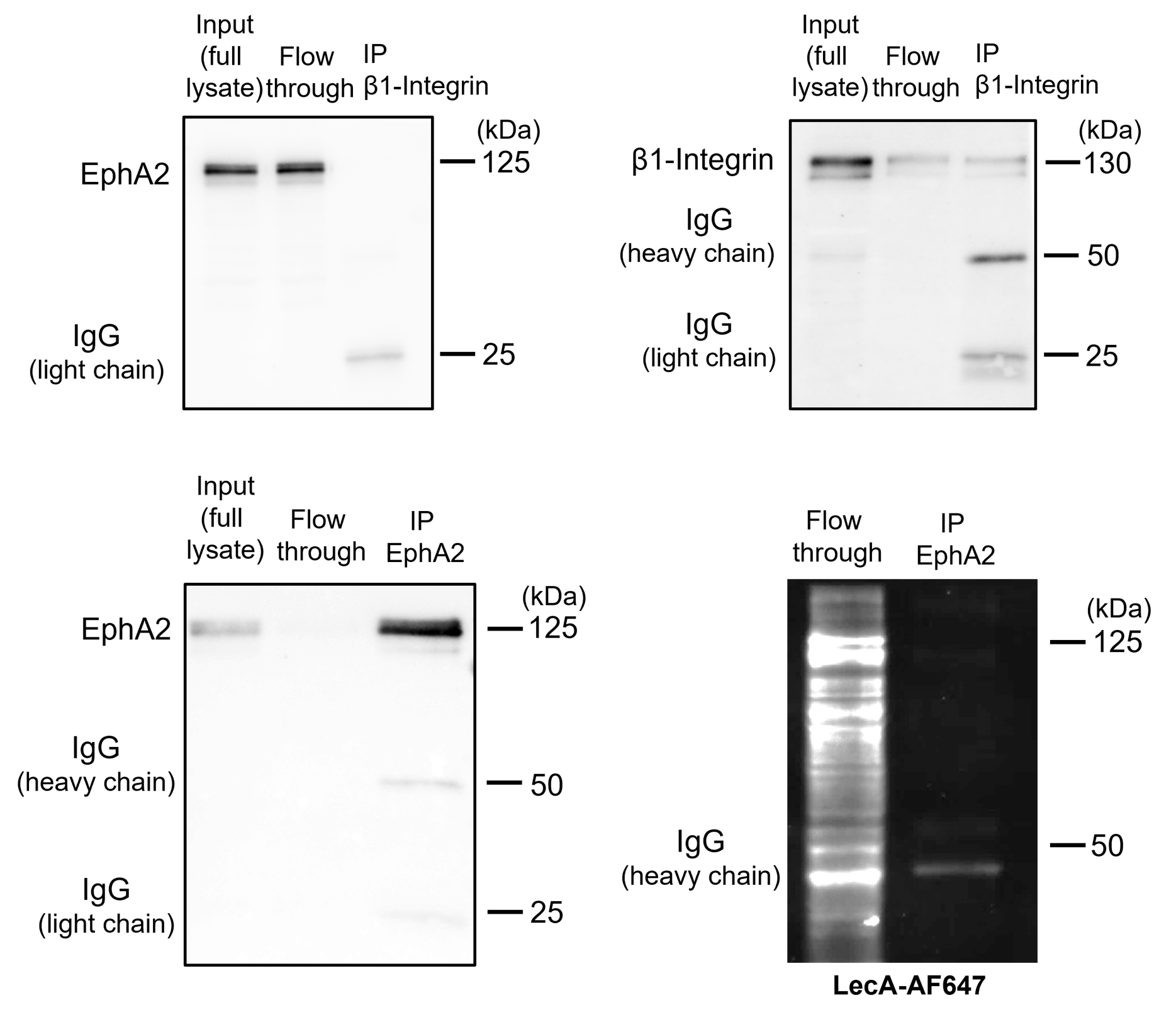

**Supplementary Fig. 6 LecA does not bind to EphA2 immunoprecipitates.**

Whole-cell lysates of H1299 cells were incubated with anti-EphA2 antibodies and immunoprecipitated using superparamagnetic Protein G Dynabeads. The whole-cell lysate (input), flow-through (unbound fraction prior to elution), and eluted antibody–antigen complexes (IP) were analyzed by immunoblotting using an anti-EphA2 primary antibody and an HRP-conjugated secondary antibody (left). In parallel, fluorescently labeled LecA-AF647 showed no binding to immunoprecipitated EphA2 in a far-Western lectin blot (right). Bands corresponding to IgG heavy and light chains result from cross-reactivity of the secondary antibody with the immunoprecipitating antibody.

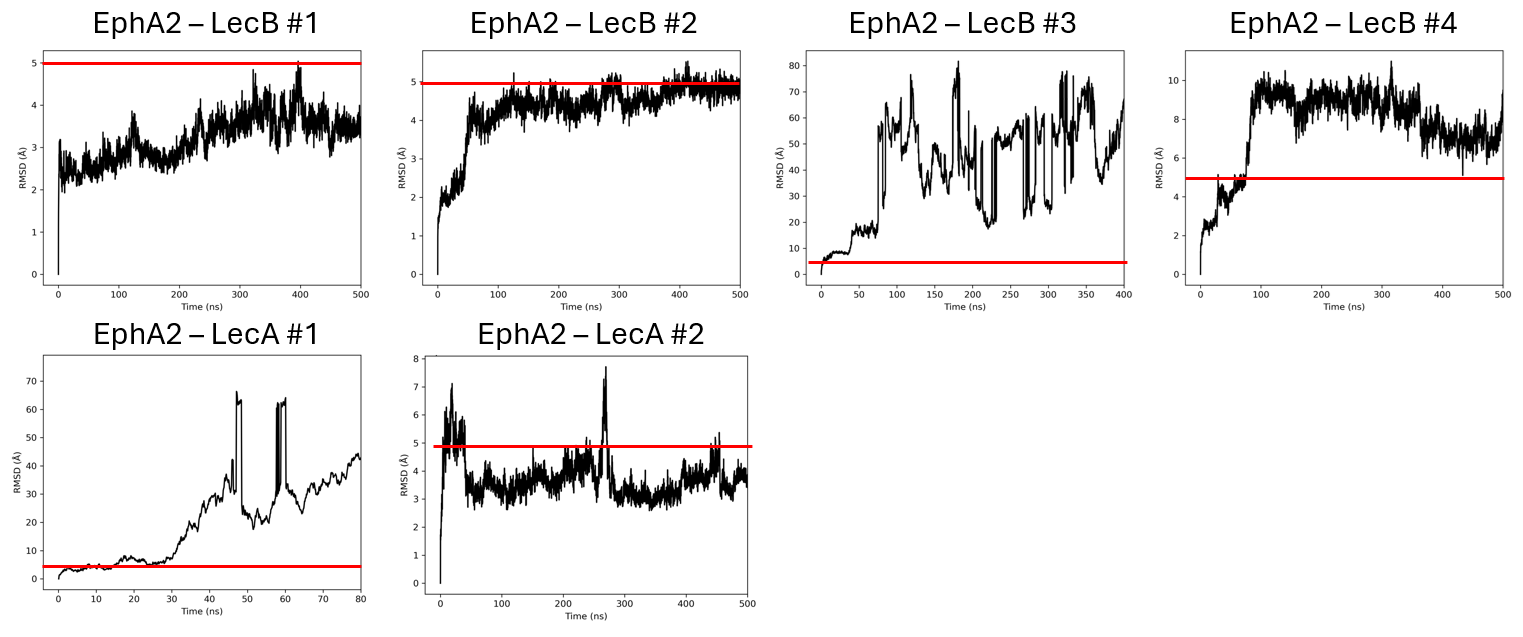

**Supplementary Fig. 7 Stability of EphA2 complexes with LecA and LecB assessed by MD simulations.**

Root-mean-square deviation (RMSD) calculated for all Cα atoms of the molecular system along the molecular dynamics (MD) trajectory, aligned to the Cα atoms of EphA2 in the initial frame. Exceeding the 5 Å threshold (red horizontal line) indicates substantial displacement of the lectin relative to EphA2, culminating in complete dissociation of the complex. In replicates exhibiting early complex dissociation (EphA2-LecB #3 and EphA2-LecA #1), simulations were terminated prior to reaching the target duration of 500 ns.

**Supplementary Table 1.** **Residues mediating stable interactions between EphA2 and LecA or LecB throughout MD simulation trajectories.**

Interacting residues located within 4 Å of the binding partner in stable complexes, calculated over the final 100 ns of the molecular dynamics (MD) trajectories, along with mean inter-residue distances (nm) and standard deviations (SD, nm). Calculated binding free energies of the complexes are reported in kcal/mol.

| Residue | Distance | Standard deviation | Residue | Distance | Standard deviation | Binding energy |
| --- | --- | --- | --- | --- | --- | --- |
| **EphA2 – LecA #2** | | | | | | |
| Eph |  |  | LecA |  |  |  |
| H333; Y334; T336; A337; V338; G339; M340; R347; T349; V432; S433; I434; N435; Q436; T437; P439; Q506; G518; S519; K520; V521 | 3.10; 2.13; 2.97; 3.29; 2.40; 2.75; 2.93; 3.25; 3.16; 3.74; 3.37; 3.78; 3.64; 2.58; 3.65; 3.15; 3.20; 3.01; 2.86; 1.73; 2.12 | 1.34; 0.56; 1.22; 0.99; 0.40; 1.15; 1.15; 1.40; 1.44; 1.30; 1.14; 1.45; 1.73; 0.84; 1.72; 0.96; 0.92; 1.15; 1.37; 0.20; 0.28 | G37_A; P38_A; T39_A; Q40_A; D100_A; V101_A; P102_A; G103_A; T104_A; Y105_A; N107_A; N108_A; P38_B; Q40_B; W42_B; E49_B; H50_B | 2.88; 2.42; 2.23; 2.40; 1.76; 1.96; 2.29; 2.38; 2.07; 3.22; 2.36; 3.68; 3.04; 2.70; 3.78; 3.53; 2.78 | 0.84; 0.46; 0.54; 0.63; 0.21; 0.29; 0.56; 0.53; 0.64; 1.04; 0.62; 0.71; 1.83; 1.08; 1.44; 1.67; 0.90 | -26.3±10.5 |

| Residue | Distance | Standard deviation | Residue | Distance | Standard deviation | Binding energy |
| --- | --- | --- | --- | --- | --- | --- |
| **EphA2 – LecB #1** | | | | | | |
| Eph |  |  | LecB |  |  |  |
| E311; G312; F313; R327; P328; P329; S330; A331; H333; Y334; L335; T336; V338; R347; T349; Q352; D353; R425; S426; R428 | 3.19; 2.57; 3.03; 2.55; 2.35; 2.92; 2.01; 3.38; 2.26; 2.32; 2.96; 2.49; 2.63; 2.60; 3.49; 2.20; 2.37; 3.64; 2.70; 2.11 | 0.98; 0.43; 0.91; 0.32; 0.23; 0.67; 0.40; 0.50; 0.32; 0.24; 0.82; 0.53; 0.80; 0.48; 0.91; 0.62; 0.57; 1.54; 0.46; 0.48 | A1_B; T2_B; Q3_B; N70_B; G71_B; R72_B; P73_B; D75_B; L76_B; V77_B; E95_B; N100_B; Y102_B; A1_D; T2_D; Q3_D; L31_D; N34_D; E35_D; Q64_D; Q66_D; P73_D; D75_D | 3.27; 3.37; 2.32; 2.69; 2.42; 2.00; 2.27; 2.03; 2.21; 2.84; 3.91; 2.87; 2.76; 3.00; 2.25; 2.71; 3.79; 3.69; 3.45; 2.90; 2.53; 2.87; 3.53 | 1.19; 1.15; 0.45; 0.76; 0.24; 0.18; 0.47; 0.47; 0.38; 0.63; 0.69; 0.65; 0.41; 0.58; 0.33; 0.38; 1.23; 1.67; 2.19; 1.13; 0.60; 0.73; 0.95 | -5.7±12.5 |

| Residue | Distance | Standard deviation | Residue | Distance | Standard deviation | Binding energy |
| --- | --- | --- | --- | --- | --- | --- |
| **EphA2 – LecB #2** | | | | | | |
| Eph |  |  | LecB |  |  |  |
| E311; G312; R327; P328; P329; S330; A331; H333; Y334; Q352; D353; C369; P371; G374; E375; C376; N407; T409; T423; S424; R425; S426; F427; R428; T429; A430; S431; S433 | 2.06; 2.81; 1.81; 2.07; 2.65; 1.99; 2.21; 2.54; 2.43; 2.32; 3.50; 3.33; 3.59; 2.41; 3.21; 3.36; 2.45; 3.92; 2.99; 1.89; 2.23; 2.33; 2.61; 2.24; 2.44; 3.00; 2.20; 2.54 | 0.44; 0.39; 0.15; 0.22; 0.21; 0.18; 0.19; 0.32; 0.21; 0.58; 0.54; 0.69; 0.98; 0.62; 0.89; 0.66; 0.44; 0.70; 0.49; 0.14; 0.27; 0.20; 0.29; 0.24; 0.29; 0.59; 0.37; 0.46 | S23_C; G24_C; T25_C; S41_C; G42_C; Q43_C; S44_C; T45_C; N46_C; N47_C; A48_C; V49_C; I50_C; G51_C; N70_C; R72_C; T25_D; F40_D; S41_D; G42_D; Q43_D; S44_D; T45_D; N46_D; A48_D; V49_D; I50_D; T52_D; Q53_D; V54_D | 2.08; 3.57; 2.51; 1.88; 2.31; 1.80; 2.35; 1.96; 2.23; 3.56; 2.40; 2.92; 3.06; 2.76; 2.46; 3.81; 3.87; 3.20; 3.96; 3.92; 2.22; 2.96; 3.38; 2.25; 2.52; 2.20; 3.07; 2.51; 2.48; 3.00 | 0.34; 0.49; 0.22; 0.13; 0.20; 0.11; 0.19; 0.19; 0.33; 0.59; 0.20; 0.41; 0.41; 0.38; 0.36; 2.05; 1.56; 0.51; 1.50; 0.72; 0.37; 0.51; 0.76; 0.40; 0.39; 0.32; 0.64; 0.26; 0.24; 0.51 | -35.6±9.2 |
